# Dot1L interacts with Zc3h10 to activate UCP1 and other thermogenic genes

**DOI:** 10.1101/2020.06.22.154963

**Authors:** Danielle Yi, Hai P. Nguyen, Jennie Dinh, Jose A. Viscarra, Ying Xie, Jon M. Dempersmier, Yuhui Wang, Hei Sook Sul

## Abstract

Brown adipose tissue is a metabolically beneficial organ capable of dissipating chemical energy into heat, thereby increasing energy expenditure. Here, we identify Dot1L, the only known H3K79 methyltransferase, as an interacting partner of Zc3h10 that transcriptionally activates the UCP1 promoter and other BAT genes. Through a direct interaction, Dot1L is recruited by Zc3h10 to the promoter regions of thermogenic genes to function as a coactivator by methylating H3K79. We also show that Dot1L is induced during brown fat cell differentiation and by cold exposure and that Dot1L and its H3K79 methyltransferase activity is required for thermogenic gene program. Furthermore, we demonstrate that Dot1L ablation in mice using UCP1-Cre prevents activation of UCP1 and other target genes to reduce thermogenic capacity and energy expenditure, promoting adiposity. Hence, Dot1L plays a critical role in the thermogenic program and may present as a future target for obesity therapeutics.

## INTRODUCTION

Adipose tissue has a central role in controlling mammalian energy metabolism in that, while white adipose tissue (WAT) is to store excess calories, brown adipose tissue (BAT) is to dissipate energy via non-shivering thermogenesis (Cannon and Nedergaard, 2004). Classic brown adipocytes contain a high density of mitochondria that contain Uncoupling protein 1 (UCP1), which uncouples respiration from ATP synthesis and generates heat instead (Farmer, 2008, Cannon and Nedergaard, 2004). In addition, beige/brite adipocytes in WAT depot are recruited upon cold exposure or β_3_-adrenergic stimulation (Seale et al., 2008, Wang and Seale, 2016, Sanchez-Gurmaches et al., 2016, Cypess et al., 2009, Lichtenbelt et al., 2009, Virtanen et al., 2009, Chondronikola et al., 2014). Based on several cross-sectional studies, adult human brown fat or brown fat-like tissue is inversely correlated with body mass index and visceral fat (Lichtenbelt et al., 2009, Hibi et al., 2016, Jimenez et al., 2007). Therefore, unraveling the mechanisms underlying the thermogenic gene program has drawn growing attention in obesity research as a promising avenue to combat obesity and associated metabolic diseases.

One of the major advances in BAT biology has been understanding the transcription network that governs the thermogenic gene program and finding critical factors that activate the UCP1 gene. A multitude of transcriptional regulators have been implicated in the transcription of UCP1, including transcription factors, Zfp516, IRF4 and EBF2, and transcriptional coregulators, PRDM16, PGC1α and LSD1 (Seale et al., 2008, Rajakumari et al., 2013, Dempersmier et al., 2015, Puigserver et al., 1998, Sambeat et al., 2016, Kong et al., 2014). Recently, we identified a BAT-enriched and cold-induced transcription factor, Zc3h10 that activates the UCP1 and other target genes, such as Tfam and Nrf1 for mitochondrial biogenesis (Yi et al., 2019). Zc3h10 activates the UCP1 promoter by directly binding to the -4.6 kb region. Upon sympathetic stimulation, Zc3h10 is phosphorylated at S126 by p38 mitogen-activated protein kinase (MAPK) to increase binding to this distal region of the UCP1 promoter. Consequently, Zc3h10 ablation in mice impairs the thermogenic gene program, while Zc3h10 overexpression in adipose tissue enhances the thermogenic capacity and energy expenditure, protecting mice from diet-induced obesity (Yi et al., 2019).

As in most biological processes, BAT as well as beige fat, rely heavily on environmental cues for its full activation of the thermogenic gene program. Integrating epigenetic effectors into the thermogenic network may provide a comprehensive understanding in the regulation of thermogenic gene program (Yi et al., 2020). Here, we identify Dot1L (disruptor of telemetric silencing 1-like) as an interacting partner of Zc3h10 and a critical coactivator of thermogenic genes. Dot1L is the only known methyltransferase that catalyzes the sequential mono-, di- and tri-methylation of H3K79, which, unlike major epigenetic sites, is located at the globular domain of nucleosome (Min et al., 2003, van Leeuwen et al., 2002, Frederiks et al., 2008). Differing from other histone methyltransferases, Dot1L does not contain a SET domain but uniquely has an AdoMET motif (Feng et al., 2002, van Leeuwen et al., 2002). Dot1L is broadly known to play roles in telomere silencing, cell cycle regulations and is particularly well studied in mixed lineage leukemia (MLL)-related leukemogenesis (Schulze et al., 2009, Okada et al., 2005, Nguyen and Zhang, 2011, Ng et al., 2002). However, not much is known about Dot1L recruitment to specific sites by specific transcription factors. Nor the role of Dot1L is thermogenic gene program is known.

We show here that Dot1L is recruited to the UCP1 promoter region via its direct interaction with Zc3h10 for Zc3h10-mediated transcriptional activation of the UCP1 and other target genes. By using the specific chemical inhibitor of Dot1L-H3K79 methyltransferase activity, pinometostat (EPZ-5676), we demonstrate that Dot1L methyltransferase activity is required for thermogenic gene expression in vitro and in vivo. Moreover, Dot1L ablation in brown adipocytes impairs, while ectopic Dot1L expression enhances, thermogenic gene program and that Dot1L requires Zc3h10 for its function in thermogenesis. Dot1L ablation in UCP1^+^ cells in mice impairs the thermogenic capacity and lowers oxygen consumption, leading to weight gain. In this regard, the GWAS database reveals multiple SNPs of Dot1L associated with a waist-hip ratio and body mass index, further supporting a potential role of Dot1L in human obesity (GWAS Central identifier: HGVPM1111, HGVPM1114).

## RESULTS

### Dot1L directly interacts with Zc3h10 for its recruitment and activation of UCP1 and other thermogenic genes

We previously reported Zc3h10 as a BAT-enriched transcription factor that promotes the BAT gene program (Yi et al., 2019). Most transcription factors do not work alone but rather form a complex to recruit other cofactors for transcription. Thus, as a DNA-binding protein, Zc3h10 may interact with coregulators to activate BAT gene transcription. Therefore, we searched for potential interacting partners of Zc3h10. The Zc3h10 with the streptavidin- and calmodulin-binding epitope-tag were incubated with nuclear extracts from BAT and the Zc3h10 complex after sequential purification were subjected to mass spectrometry. We identified multiple potential Zc3h10 interacting proteins, among which were methyltransferases and chromatin remodelers (Figure S1A). For further studying as a Zc3h10 interacting protein, we selected Dot1L (disruptor of telemetric silencing 1-like), the H3K79 methyltransferase, as it directly interacted with Zc3h10 and was the only BAT enriched gene out of the candidates tested.

First, for validation of interaction between Dot1L and Zc3h10, we performed Co-IP experiments with lysates of HEK293FT cells transfected with HA-Dot1L and Flag-Zc3h10 by using FLAG and HA antibodies. Indeed, we detected FLAG-Zc3h10 upon immunoprecipitation with HA antibodies. Conversely, HA-Dot1L was detected upon immunoprecipitation with FLAG antibodies (Figure 1A, left). We then tested an interaction of the two endogenous proteins using BAT from mice. Again, we detected the presence of endogenous Zc3h10 when nuclear extracts of BAT from mice were immunoprecipitated with Dot1L antibody. By reverse Co-IP, we confirmed the interaction between endogenous Dot1L and Zc3h10 (Figure 1A, right). We next asked whether Dot1L can directly bind Zc3h10, by using glutathione S-transferase (GST) fused to Dot1L expressed and purified from *E. Coli* (Figure S1B). Incubation of GST-Dot1L fusion protein immobilized on glutathione beads with *in vitro* transcribed and translated [^35^S]-Zc3h10, but not control GST-alone, detected the presence of Zc3h10 (Figure 1B). Overall, these results demonstrate that Dot1L directly interacts with Zc3h10. We next examined domains of Dot1L for its interaction with Zc3h10. We generated three Dot1L truncated constructs; N-terminal Dot1L (1-500aa) containing the catalytic domain, the middle fragment (501-1000aa) containing the coiled-coil domain, and the C-terminal domain (1001-1540aa); All constructs were N-terminally tagged with HA. We then cotransfected these constructs with full-length Flag-tagged Zc3h10. By Co-IP, we detected the middle fragment (501 -1000) of Dot1L, that contains coiled coil motifs, interacting with Zc3h10, but not the other two regions of Dot1L (Figure 1C). We conclude that the middle fragments of Dot1L contains the Zc3h10 interacting domain.

**Figure 1.**
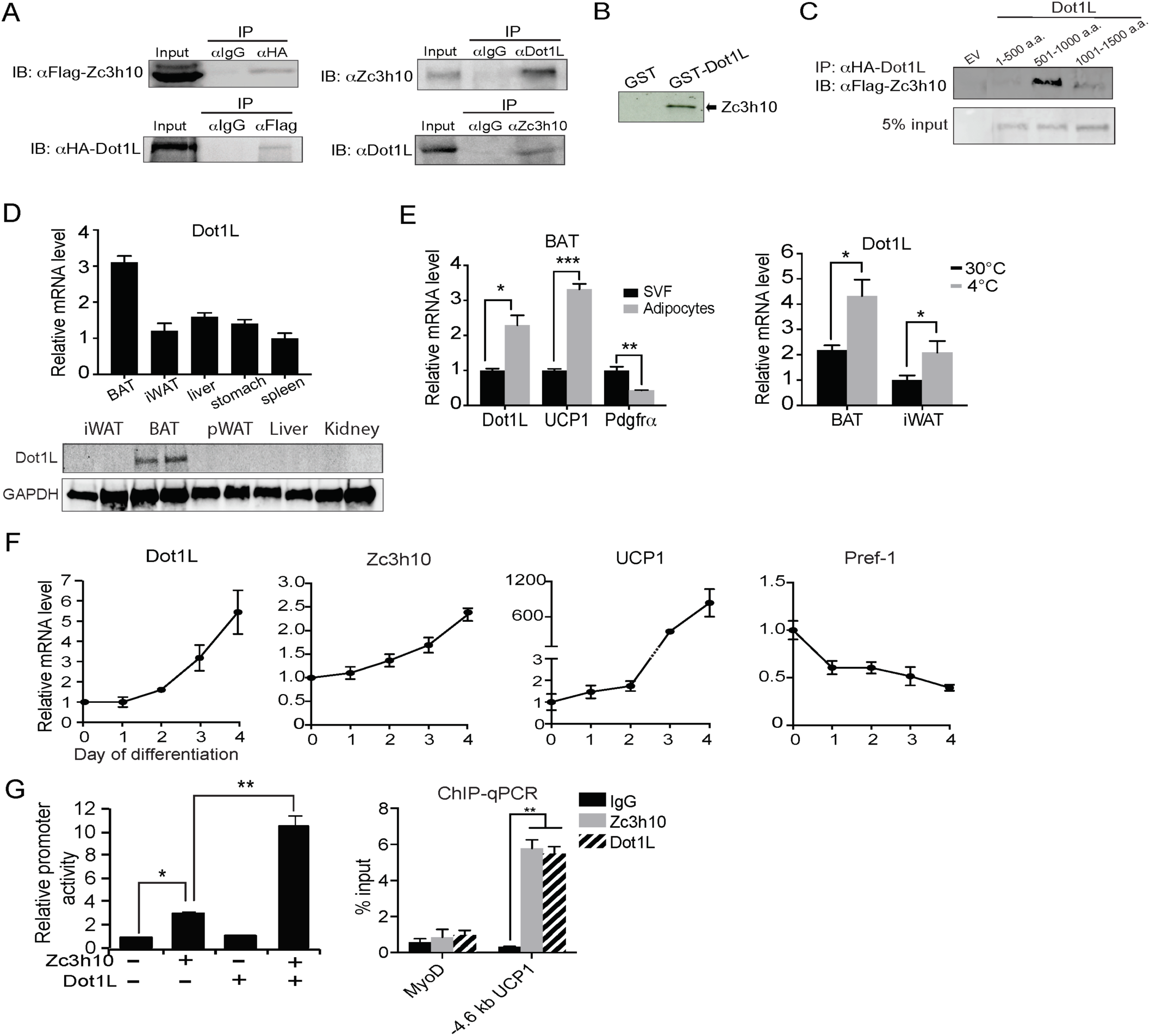
DotIL directly interacts with Zc3h10 for its recruitment and activation of BAT gene program. (A) (Left) ColP using αFlag for Zc3h10 or αHA for DotIL after immunoprecipitation with either αHA or αFlag, respectively, using lysates from HEK293FT cells transfected with Flag-Zc3h10 and HA-Dot1 L. (Right) ColP of endogenous Zc3h10 and Dot1 L protein using BAT tissue of C57BL/6 using αZc3h10 and αDotlL. (B) Autoradiograph of GST pull-down using GST-Dot1 L and 35S-labeled in vitro transcribed/ translated Zc3h10. (C) ColP using αFlag for Zc3h10 after immunoprecipitation with αHA for Dot1 L using lysates from HEK293FT cells transfected with Flag-Zc3h10 and various HA-Dot1 L constructs. (D) RT-qPCR and western blot analysis of Dot1 L in various tissues from 10-week-old C57BL/6 mice (n=5). (E) (Left) RT-qPCR for indicated genes in the adipocyte fraction and SVF from BAT. (Right) RT-qPCR for Dot1 L mRNA of BAT and iWAT from mice housed at either 30°C or4°C (n=5). (F) RT-qPCR for indicated genes during the course of BAT cell differentiation. (G) (Left) HEK293FT cells were cotransfected with the -5.5kb UCP1-Luc promoter with Zc3h10 or Dot1 L either together or individually (n=5). (Right) ChlP-qPCR for Zc3h10 and Dot1 L enrichment at the -4.6kb region of UCP1 promoter using Flag or HA antibodies. Differentiated BAT cells were transduced with either Flag-Zc3h10 or HA-Dot1 L. Data are expressed as means ± standard errors of the means (SEM). *p < 0.05, **p < 0.01, ***p < 0.001. Supplementary Figure S1 is linked to Figure 1.

Since Zc3h10 activates UCP1 and other target genes for BAT gene program and Dot1L interacts with Zc3h10, Dot1L expression pattern could be similar to Zc3h10 and thus BAT-enriched. Indeed, tissue expression profiling by RT-qPCR and immunoblotting showed that Dot1L was enriched in mouse BAT compared to other tissues tested, such as WAT, liver and stomach (Figure 1D). We next compared Dot1L mRNA levels between the stromal vascular fraction (SVF) that contains preadipocytes with adipocyte fraction of BAT from 10-wk-old wild-type (WT) mice. As expected, UCP1 was enriched in the adipocyte fraction, whereas PDGFRα expression was higher in the SVF fraction. We found Dot1L to be enriched in the brown adipocyte fraction by over 2-fold compared to the SVF (Figure 1E, left). Moreover, similar to Zc3h10, expression of Dot1L in BAT was induced upon cold exposure (Figure 1E, right). We next examined Dot1L expression during BAT cell differentiation in vitro. During the course of brown adipocyte differentiation, as expected, UCP1 expression was induced, whereas expression of preadipocyte gene, Pref-1, was suppressed. More importantly, similar to Zc3h10, Dot1L mRNA level was increased during BAT cell differentiation (Figure 1F). Overall, we conclude that Dot1L expression pattern is similar to Zc3h10 in terms of enrichment in mature brown adipocytes and induction by cold exposure, which may allow potential a cooperative function of Dot1L and Zc3h10 for the BAT gene program.

Next, to examine the functional significance of Dot1L and Zc3h10 interaction in the activation of UCP1 promoter, we performed the luciferase (Luc) reporter assay using the -5.5 kb UCP1-Luc promoter. Along with the UCP1 promoter-Luc construct, Zc3h10 and Dot1L were cotransfected into HEK293 cells. As expected, compared to empty vector control, Zc3h10 alone activated the UCP1 promoter over 3-fold. Dot1L alone could not activate the UCP1 promoter.

When co-transfected with Zc3h10, Dot1L was able to synergistically activate the UCP1 promoter, resulting in a robust 11-fold increase in the UCP1 promoter activity (Figure 1G, left). These results demonstrate the cooperative function of Dot1L and Zc3h10 in UCP1 promoter activation. Next, because Dot1L interacts with Zc3h10 to enhance the UCP1 promoter activity, we predicted that Dot1L should occupy the same UCP1 promoter region where Zc3h10 binds. We performed chromatin immunoprecipitation (ChIP) using BAT cells transduced with adenovirus containing Flag-tagged Zc3h10 and HA-tagged Dot1L. By using Flag (for Zc3h10) and HA (for Dot1L) antibodies, we detected strong enrichment of both Zc3h10 and Dot1L at the - 4.6 kb region of the UCP1 promoter, a region corresponding to the Zc3h10 binding site (Yi et al., 2019) (Figure 1G, right). This illustrates co-occupancy of Dot1L and Zc3h10 at the same region of the UCP1 promoter. Collectively, we demonstrate that Dot1L, as an interacting partner of Zc3h10, is a critical coactivator of the UCP1 promoter.

### Dot1L is critical for the thermogenic gene program, and its action is dependent on Zc3h10

We next tested whether Dot1L is required for BAT gene program in cultured BAT cells. BAT cells at Day 2 of differentiation were transduced with adenovirus expressing short hairpin RNAs targeting Dot1L for knockdown (Dot1L KD). Transduction of shDot1L adenovirus in BAT cells caused a decrease in Dot1L mRNA levels by approximately 60% (Figure 2A, left). While there were no apparent changes in expression of transcription factors critical for brown adipocyte differentiation, such as PPARγ, there was a 50% reduction in UCP1 expression at mRNA and protein levels (Figure 2A). Expression of other Zc3h10 target genes, including Tfam, Nrf1 and Elovl3, was also decreased significantly (Figure 2A, middle). Moreover, Dot1L KD BAT cells had decreased oxygen consumption rate (OCR) in oligomycin-, FCCP-, and rotenone/antimycin A-treated conditions (Figure 2B, left). Notably, the uncoupled respiration of Dot1L KD cells was significantly lower than the control BAT cells in both basal and in forskolin-stimulated conditions (Figure 2B, right). These results show that Dot1L is critical for full activation of the BAT gene program and mitochondriogenesis, and thus thermogenic function in BAT cells.

**Figure 2.**
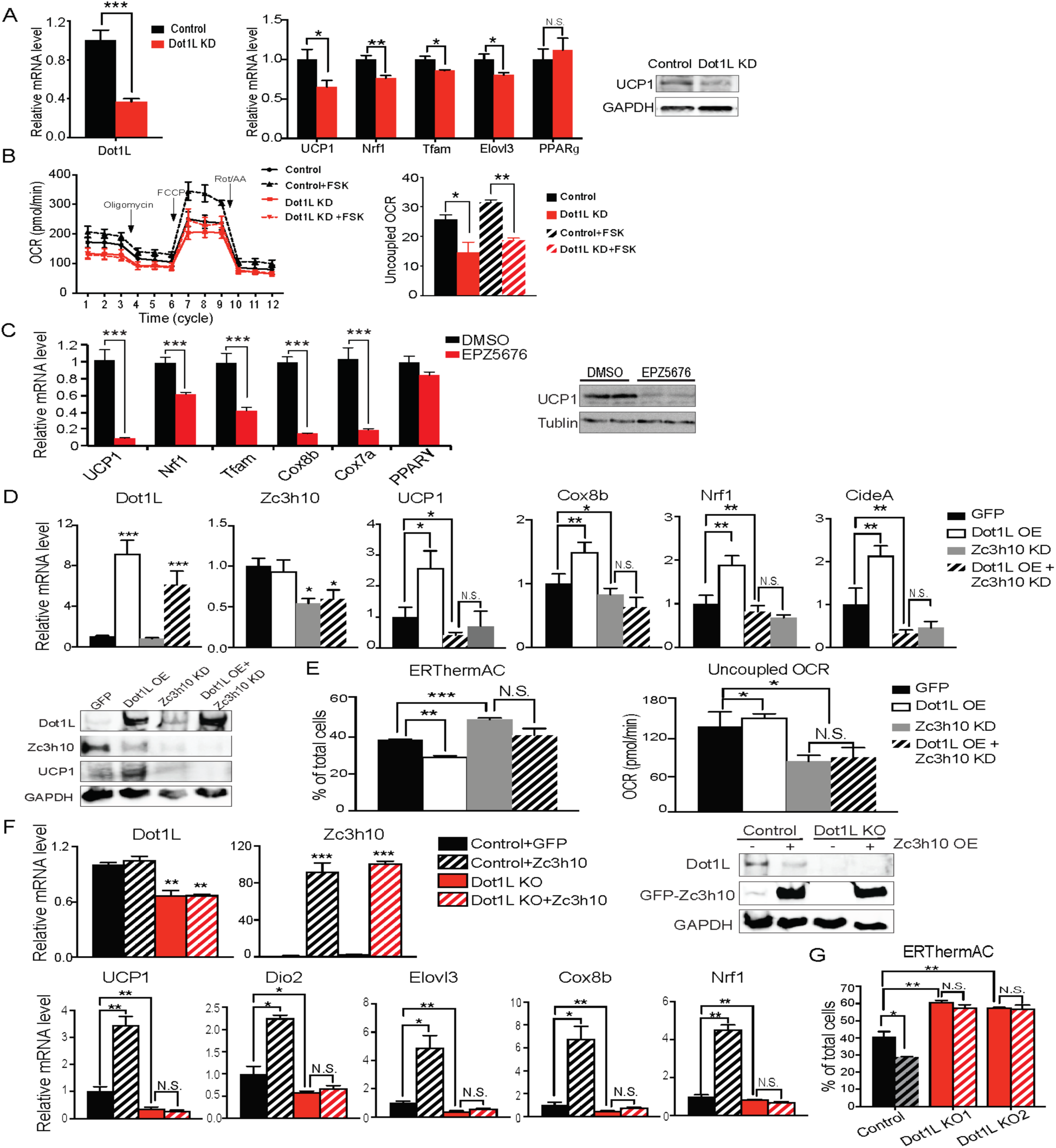
Dot1 L is critical for the thermogenic gene program, and its action is dependent on Zc3h10. (A) (Left) RT-qPCR for Dot1 L and thermogenic genes in BAT cells infected either scrambled (Control) or adenovirus expressing short hairpin targeting DotIL (DotIL KD) after D2 of adipogenic differentiation (n=6). (Right) Immunoblotting for UCP1. (B) (Left) OCR measured in DotIL KD cells using SeahorseXF24 analyzer (n=5). (Right) Uncoupled OCR in BAT cells infected with control or shDotIL under oligomycin (0.5uM). (C) (Left) RT-qPCR for indicated genes in BAT cells treated with DotIL chemical inhibitor, EPZ5676 (5nM). (Right) Western blotting analysis for UCP1 protein. (D) (Top) RT-qPCR for indicated genes and (Bottom) immunoblotting for indicated proteins in differentiated BAT cells that were transduced with either AdGFP or AdDotl L or shZc3h10 individually or in combination for overexpression of Dot1 L and knockdown of Zc3h10 (n=6). The differentiated cells were treated with forskolin (10uM) for 6 hr. (E) (Left) FACS analysis and quantification of ERthermAC, that inversely correlates with heat production. (Right) Uncoupled OCR measured in BAT cells by Seahorse assay (n=5). (F) RT-qPCR for indicated genes and immunoblotting for indicated proteins in the control BAT cells or DotlL-CRISPR KO pool, overexpressing either GFP or Zc3h10. (G) FACS analysis and quantification of ERthermAC in Zc3h10 overexpressing in the control or Dot1 L-CRISPR KO pools treated with forskolin(10uM). Data are expressed as means ± standard errors of the means (SEM). *p < 0.05, **p < 0.01, ***p < 0.001. Supplementary Figure S2 is linked to Figure 2.

Next, in order to further establish the role for Dot1L in BAT gene program, we treated BAT cells with a specific inhibitor of Dot1L H3K79 methyltransferase, EPZ5676. We predicted that if Dot1L is required for the transcriptional activation of UCP1 and other BAT genes, inhibition of Dot1L activity should prevent BAT gene expression. Dot1L inhibition resulted in a significant reduction in mRNA and protein levels for UCP1 by 90% (Figure 2C). Expression of other BAT-enriched Zc3h10 target genes, such as Tfam, Nrf1, Cox7a and Cox8b, was decreased by 40-80%, whereas PPARγ expression was not affected (Figure 2C). These results show that Dot1L methyltransferase activity is required for thermogenic gene expression and other Zc3h10 target genes for the BAT gene program.

To investigate whether Dot1L’s function is dependent on its recruitment by Zc3h10 to thermogenic genes, we performed adenoviral overexpression of Dot1L (Dot1L OE) and knockdown of Zc3h10 (Zc3h10 KD) alone or in combination in differentiated BAT cells. We verified that Dot1L was overexpressed by over 6-fold, and Zc3h10 was knocked down by 50% at the mRNA levels and protein levels (Figure 2D). As expected, Dot1L overexpression significantly increased thermogenic gene expression including UCP1, Cox8b, CideA, while Zc3h10 KD significantly decreased these genes (Figure 2D). To examine the functional changes of thermogenic capacity in these BAT cells, we utilized a small molecule thermosensitive fluorescent dye, ERthermAC in live cells (Kriszt et al., 2017). Corresponding to an increase in temperature, ERthermAC can accumulate in the ER to decrease fluorescence and, thus, fluorescence and thermogenesis are inversely correlated. Upon forskolin treatment, the population of ERthermAC^+^ cells in Dot1L OE cells was decreased by 10%, which indicated that the cell population with higher temperature was increased by approximately 2.5-fold (Figure 2E, Left), calculated by the percent decrease from ERthermAC^+^ population in control cells. In contrast, ERthermAC^+^ cells in Zc3h10 KD cells were significantly increased by 10%, indicating that Zc3h10 ablation decreased heat production in BAT cells. Moreover, Zc3h10 KD in Dot1L overexpressing cells showed low ERthermAC^+^ cells that were similar to control cells, indicating that Zc3h10 ablation prevented Dot1L-induced thermogenesis. Furthermore, when we measured OCR by using Seahorse, Dot1L overexpression in Zc3h10 KD BAT cells had the uncoupled OCR comparable to that of Zc3h10 KD, which was significantly lower than the control cells (Figure 2E, Right). These results are in accord with the concept that Dot1L activation of thermogenic genes is dependent on Zc3h10.

Next, we asked whether Dot1L is required for Zc3h10-induced thermogenesis. To induce Dot1L ablation in vitro, we used Dot1L knockout (Dot1L KO) pools generated by the CRISPR inducible Cas9 system. We then overexpressed Zc3h10 in both control and the Dot1L KO pool and differentiated them. We verified that Dot1L was reduced at the mRNA levels and protein levels by approximately 50% in the Dot1L KO pool, and Zc3h10 was overexpressed by more than 80-fold. We found that Dot1L KO pool had significantly decreased UCP1, Dio2, Elov3, Cox8b, and Nrf1 mRNA levels by more than 50% compared to the control BAT cells, while Zc3h10 overexpression increased thermogenic gene expression as expected (Figure 2F). Importantly, Zc3h10 overexpression in Dot1L KO pool did not rescue the decreased BAT-gene expression, remaining in significantly reduced thermogenic gene expression, an evidence that Dot1L is critical for Zc3h10-induced thermogenic gene program. We also utilized ERthermAC to test heat generation in these cells. Zc3h10 OE significantly reduced ERthermAC^+^ population in compared to control; while two independent Dot1L KO pools had increased ERthermAC^+^ population, indicating decreased thermogenesis (Figure 2G). Notably, Zc3h10 OE did not affect ERthermAC^+^ population in Dot1L KO pools. Altogether, these data further show that Dot1L is required for activation of thermogenic genes by Zc3h10.

We then tested the ability of Dot1L to induce UCP1 and other thermogenic genes by gain of function experiment using 3T3-L1 cells. We overexpressed Dot1L or Zc3h10 alone or together via adenoviral transduction in 3T3-L1 cells and the cells were induced to browning by forskolin treatment. We first verified Zc3h10 and Dot1L overexpression by RT-qPCR and immunoblotting (Figure S2A). Interestingly, Dot1L overexpression alone increased UCP1 mRNA levels over 2-fold, probably due to the presence of endogenous Dot1L in 3T3-L1 cells. Co-overexpression of both Dot1L and Zc3h10 resulted in a further increase in UCP1 mRNA levels by 3-fold and UCP1 protein levels (Figure S2A and B). Furthermore, overexpression of Dot1L and Zc3h10 together in differentiated 3T3-L1 cells further increased mRNA levels for Tfam and Nrf1 in comparison to Dot1L or Zc3h10 alone. Expression of a terminal adipose gene, FABP4, was not significantly different in all conditions. With increased expression of UCP1 and other BAT-enriched genes upon overexpression of Dot1L and Zc3h10 (Figure S2A), we next assessed the functional consequence of the oxygen consumption. Indeed, forskolin-treated Dot1L alone overexpressing cells had somewhat increased OCR. Co-overexpression of both Dot1L and Zc3h10 further increased the total OCR as well as uncoupled OCR, suggesting cooperative function of Dot1L and Zc3h10 in browning of 3T3-L1 cells (Figure S2C). Altogether, these results demonstrate that Dot1L plays a critical role in promoting thermogenic program.

### Inhibition of Dot1L methyltransferase activity impairs BAT gene program in vivo

To study Dot1L function in vivo, we first used a selective and potent small molecule inhibitor of Dot1L, EPZ5676. We administered either saline (control) or EPZ5676 to WT mice via daily subcutaneous injection for 8 days (Figure 3A). By immunoblotting with H3K79me3 antibody, we confirmed inhibition of H3K79 methylation in BAT of animals treated with EPZ5676 (Figure 3B, left). Indeed, BAT from the inhibitor treated mice showed significantly decreased mRNA levels for UCP1, Nrf1, and Tfam, as well as other thermogenic genes, such as PGC1α and Elovl3, while no changes in adipogenic transcription factor, such as PPARγ (Figure 3B, middle). Also, the immunoblotting detected significantly lower UCP1 protein levels in BAT of EPZ5676 treated mice compared to the control mice (Figure 3B, middle). Similarly, EPZ5676 treatment also decreased thermogenic gene expression in iWAT, WAT depot known to undergo browning (Figure 3B, right), but not in pWAT or in liver (Figure S3A).

**Figure 3.**
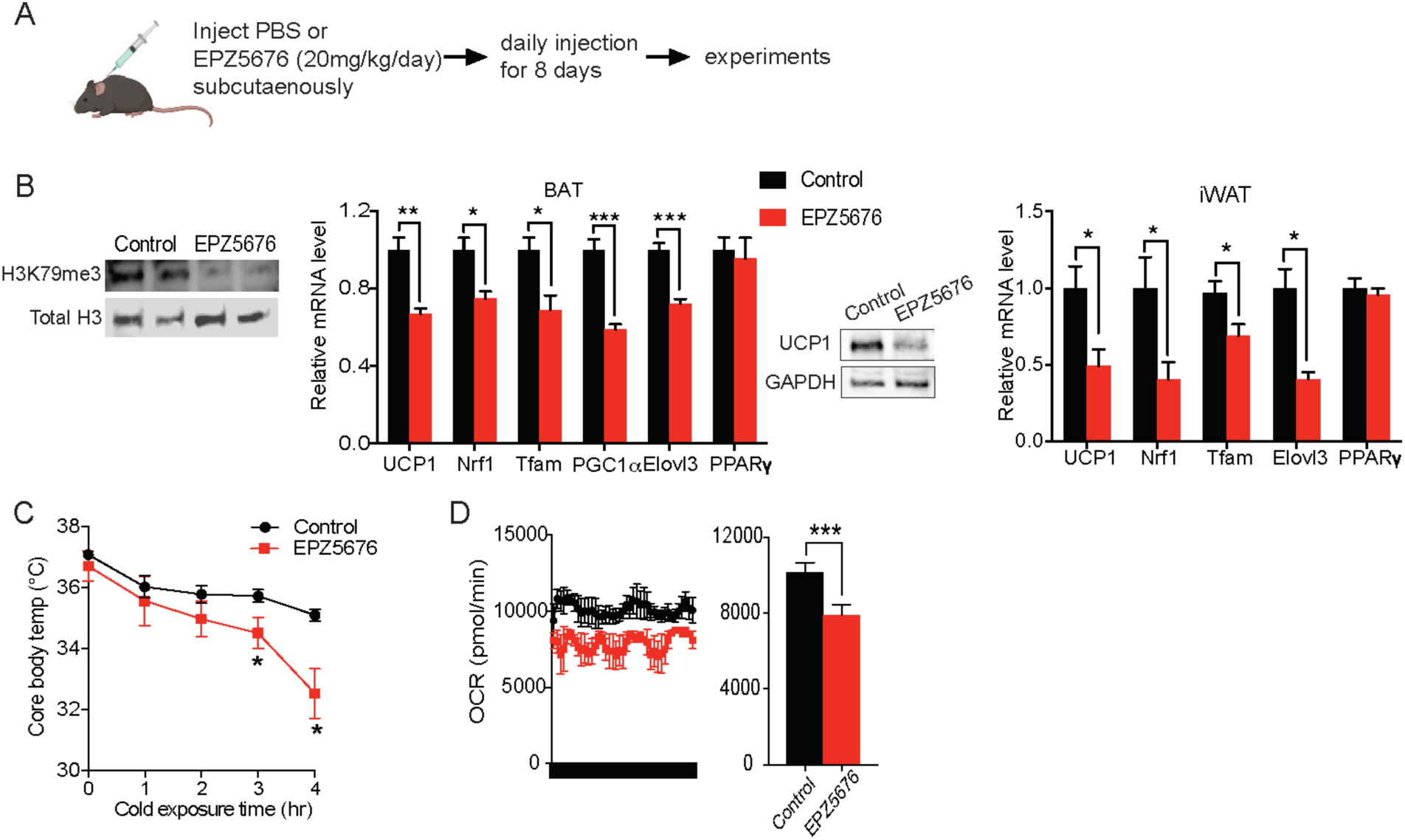
Inhibition of DotIL activity impairs BAT gene program in vivo. (A) Schematic diagram of the strategy used to inject either Saline for control or Dot1 L chemical inhibitor, EPZ5676. (B) (Left) Immunoblotting for H3K79me3 and total H3. (Middle) RT-qPCR for indicated genes in BAT and iWAT from either PBS or EPZ5676 injected mice (n=4 per group) and immunoblotting for UCP1 protein. (C) Rectal temperature of 14-wk old mice maintained at 4’C at indicated time points (hr) (n=4 per group). (D) VO2 assayed in mice that were housed at 4°C by indirect calorimetry using CLAMS. Data are expressed as means ± standard errors of the means (SEM). *p < 0.05, **p < 0.01, ***p < 0.001. Supplementary Figure S3 is linked to Figure 3.

Next, to investigate the physiological outcome from decreased expression of thermogenic genes, we subjected these EPZ5676 treated mice to acute cold exposure at 4°C. Both groups of mice had body temperatures around 37°C prior to the cold exposure. After 4 hrs of cold exposure, the body temperature of the EPZ5676 treated mice was 4°C lower compared to their non-treated littermates, demonstrating the significantly reduced thermogenic capacity (Figure 3C). Next, we assessed the energy expenditure by measuring the whole body O_2_ consumption using CLAMS. Indeed, EPZ5676 treated mice had significantly lower VO_2_ than the control mice at 4°C (Figure 3D), whereas food intake and locomotive activity were similar between the two groups (Figure S3B). Altogether, these results support that Dot1L enzymatic activity is critical for activating the thermogenic gene program in vivo.

### The requirement of Dot1L for cold-induced thermogenesis and the Dot1L-Zc3h10 function in mice

To further evaluate whether Dot1L is critical for the BAT gene program in vivo, we performed Dot1L ablation in UCP1^+^ cells in mice (Dot1L-BKO) by crossing Dot1L floxed mice with UCP1-Cre mice (Figure 4A, left). We validated our mouse model by genotyping and by comparing Dot1L mRNA levels in BAT and iWAT of Dot1L-BKO mice and Dot1L f/f control mice (WT). Dot1L-BKO mice had a 70% reduction in Dot1L mRNA level in BAT. Dot1L-BKO mice also showed a 60% decrease in Dot1L mRNA level in iWAT, probably due to the mild cold exposure at room temperature that can induce UCP1^+^ adipocytes in iWAT (Figure 4A, right). Importantly, UCP1 expression was significantly reduced by 40% and 60% in BAT and iWAT, respectively (Figure 4B). UCP1 protein level also was reduced significantly, as detect in BAT (Figure 4B right). In addition, expression of other BAT enriched genes, such as Dio2 and CideA, and other Zc3h10 target genes, such as Tfam and Nrf1, were all significantly decreased in BAT and iWAT of Dot1L-BKO mice, compared to WT littermates (Figure 4C).

**Figure 4.**
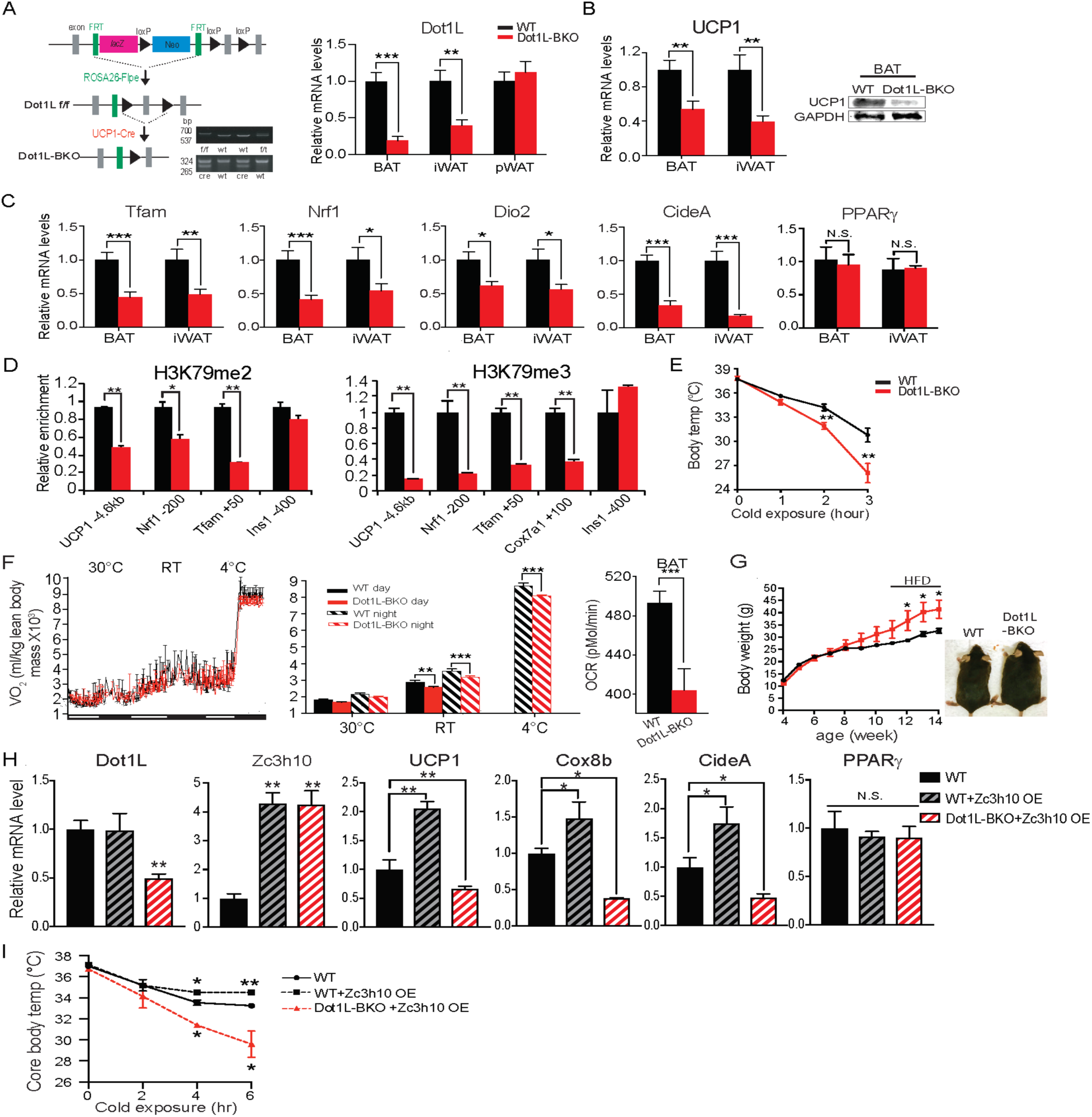
The requirement of DotIL for cold-induced thermogenesis and the Dot1 L-Zc3h 10 function in mice. (A) (Left) Schematic diagram of the strategy used to generate BAT specific Dot1 L conditional knockout mice. PCR genotyping of the mice: Top gel, DotIL allele; bottom gel, Cre. (Right) RT-qPCR for Dot1 L in BAT, iWAT and pWAT from Dot1 L f/f (WT) and Dot1L-BKO mice (n=6 per group). (B) RT-qPCR and immunoblotting for UCP1 in BAT and iWAT from Dot1 Lf/f (WT) and Dot1 L-BKO mice. (C) RT-qPCR for indicated genes in BAT and iWAT from Dot1 L f/f (WT) and Dot1 L-BKO mice. (D) ChlP-qPCR of H3K79me2 and H3K79me3 at Zc3h10 binding regions of BAT tissue from control (f/f) or Dot1 L-BKO mice (n=5 per group). (E) Rectal temperature measured in 13wk-old mice at 4°C at indicated time points (hours) (n=6 mice per group). (F) (Left) whole body V02 assayed in WT and Dot1 L-BKO mice, housed at indicated ambient temperatures (n=6 per group) by indirect calorimetry using CLAMS. (Right) OCR measured in BAT of WT and Dot1 L-BKO mice using Seahorse XF24 Analyzer (n=5 per group). (G) Representative photograph of 14wk-old control and body weight of WT and Dot1 L-BKO mice fed HFD from wk 12. (H) RT-qPCR for indicated genes from WT injected with GFP or Zc3h10 adenovirus or Dot1 L-BKO mice injected with Zc3h10 adenovirus for overexpression (n=5 per group). (I) Rectal temperature measured at 4°C at indicated time points (hours) (n=5 per group). Data are expressed as means ± standard errors of the means (SEM). *p < 0.05, **p < 0.01, ***p < 0.001. Supplementary Figure S4 is linked to Figure 4.

Dot1L is known to be the only known H3K79 methyltransferase (VanLeeuwen, 2002). Hence, we performed ChIP-qPCR, to assess methylation status of H3K79 methylation at the Zc3h10 binding region of the UCP1 promoter using BAT of mice. Indeed, we detected a 50% reduction in di-methylation and a 70% reduction in tri-methylation of H3K79 at the -4.6 kb UCP1 promoter region (Figure 4D). Thus, the decreased UCP1 mRNA level upon Dot1L ablation in BAT was correlated with decreased di-methylation and tri-methylation H3K79 at the UCP1 promoter region. Similar to the -4.6 kb UCP1 promoter region, H3K79 methylation status at the - 200 bp Nrf1 promoter region and at the +50 bp Tfam region was also decreased (Figure 4D).

We previous reported these regions to contain Zc3h10 binding sites. The reduced H3K79 methylation at Zc3h10 binding sites in BAT of Dot1L-BKO mice suggest that the recruitment of Dot1L by Zc3h10 to the target genes is important for BAT gene expression. Moreover, these observations support the notion that Dot1L participates in the transcriptional activation of UCP1 and other Zc3h0 target genes by modifying H3K9 methylation status.

To examine the effect of Dot1L deficiency on the thermogenic capacity, we subjected these Dot1L-BKO mice to an acute cold exposure at 4°C. While body temperatures of mice were similar initially, Dot1L-BKO mice were severely cold intolerant as their body temperature started to drop significantly after 2 hrs of cold challenge. After 3 hrs of cold exposure, the body temperature of Dot1L-BKO mice was 5°C lower than that of control mice (Figure 4E), an in vivo evidence of the requirement of Dot1L for cold-induced thermogenesis. Considering such lower thermogenic gene expression and decreased thermogenic capacity, we assessed the metabolic effect in the Dot1L-BKO mice by measuring whole body OCR using CLAMS. Indeed, the Dot1L-BKO mice had significantly reduced VO_2_ compared to WT littermates at room temperature and at 4°C in particular, while locomotor activity and food consumption were similar (Figure 4F and Figure S4A). To examine the contribution of BAT to the altered energy expenditure in Dot1L-BKO mice, by Seahorse assay, we measured OCR in BAT dissected out from these mice. We found OCR in BAT from Dot1L-BKO mice to be decreased by 20%. (Figure 4F, right). In line with these results, we then asked whether decreased energy expenditure and impaired BAT function in Dot1L-BKO mice would be reflected in changes in adiposity. The body weights of mice started to diverge starting on wk 10, then they were maintained on a high-fat-diet (HFD) from wk 11 to wk 13 of age. By 13 wks, the Dot1L-BKO mice were approximately 8 g heavier relative to the control mice (Figure 4G), and the Dot1L-BKO had significantly larger iWAT and pWAT depots, which primarily accounted for the higher total body weights of Dot1L-BKO mice (Figure S4B). There were no differences in food/energy intake between the two genotypes despite differences in body weights. In addition, blood glucose levels of Dot1L-BKO mice on HFD were significantly higher during the course of glucose tolerance test (GTT) (Figure S4C Left), and they had significantly impaired insulin sensitivity when subjected to an insulin tolerance test (ITT) (Figure S4C, Rght). Collectively, these results establish a defective BAT function and impaired thermogenic capacity that affects adiposity and consequently insulin sensitivity in Dot1L-BKO mice and that Dot1L is required for the BAT thermogenic program in vivo.

Next, to test the in vivo relevance of Dot1L-Zc3h10 interaction to thermogenesis, we overexpressed Zc3h10 in WT and in Dot1L-BKO mice by direct injection of Zc3h10 adenovirus into BAT. We detected a 4-fold increase in Zc3h10 mRNA levels in BAT of Zc3h10 adenovirus-injected mice (Zc3h10 OE). As expected, Dot1L-BKO mice showed a 60% reduction in Dot1L mRNA levels in BAT (Figure 4H). As we previously have reported that overexpression Zc3h10 in adipose tissue enhances thermogenic gene expression (Yi et al., 2019), Zc3h10 overexpressing mice by injection had increased expression of BAT-enriched genes, such as UCP1, Cox8b, and CideA, compared to the control mice. More importantly, expression BAT-enriched genes remained significantly reduced in Dot1L-BKO even upon Zc3h10 overexpression, while adipogenic markers, such as PPARγ, remained unchanged (Figure 4H). Moreover, when these mice were subjected to cold exposure, Zc3h10 OE mice had higher core body temperature compared to the control mice after 4 hrs (Figure 5I). However, Dot1L-BKO Zc3h10 OE mice were severely cold sensitive as their body temperature was significantly lower than the control and Zc3h10 OE mice. Taken together, these results demonstrate the requirement of Dot1L-Zc3h10 interaction, both in vitro and in vivo for activation of the thermogenic program in brown adipose tissue.

**Figure 5.**
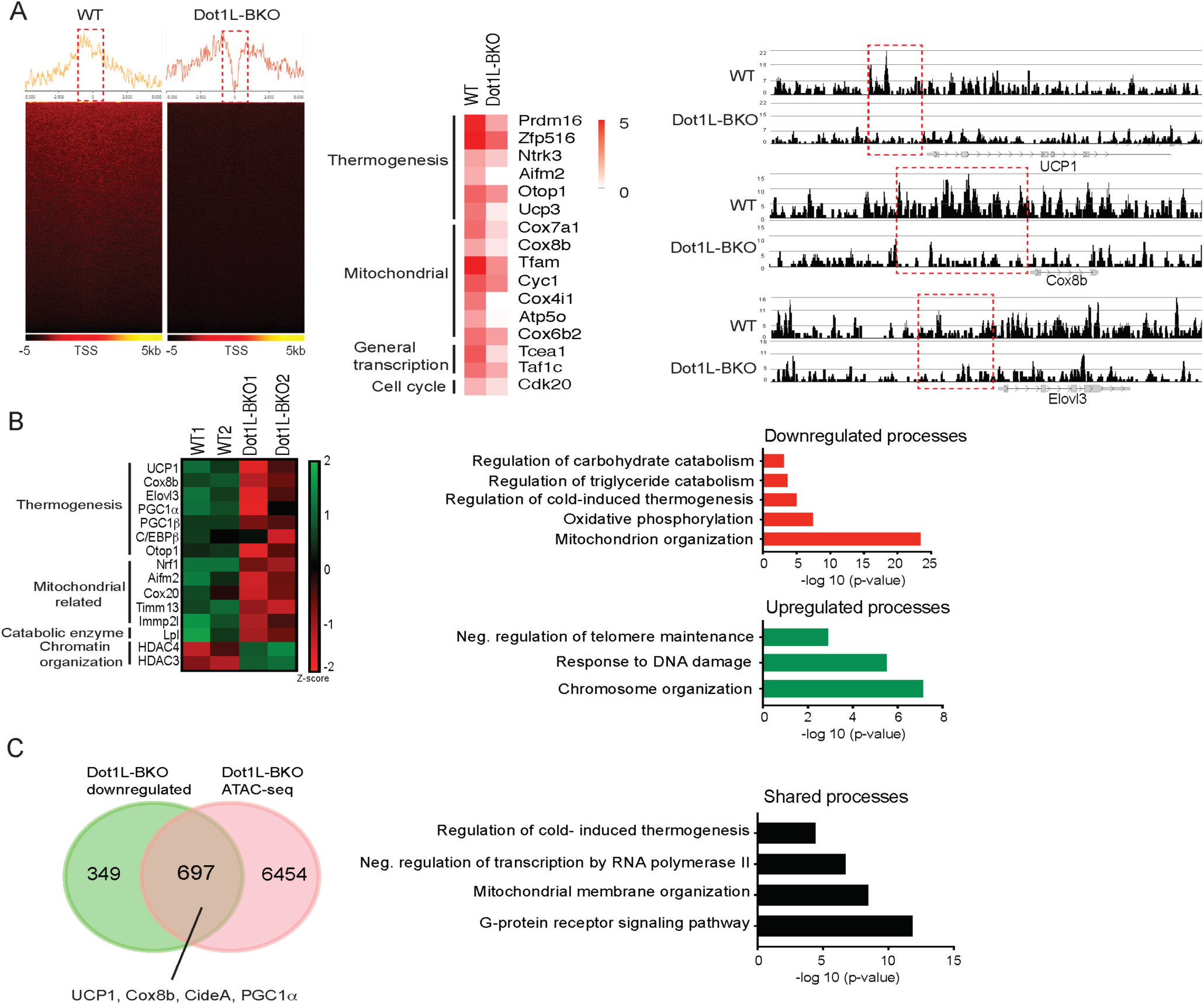
Genome wide analysis for Dot1 L effect on chromatin accessibility for thermogenic gene program. (A) ATAC-seq using BAT from WT and Dot1 L-BKO mice after 2 hr cold exposure n = 2 per group. (Left) Heatmaps showing open chromatin regions focused at the transcription start site (TSS). Color scale shows peaks detected. (Right) UCSC genome browser screenshot of representative peaks at a subset of promoter regions of thermogenic genes. (B) RNA-seq using BAT from WT and Dot1 L-BKO mice, n = 2 pooled RNA samples per group. (Left) Heatmap showing changes in gene expression. Color scale shows changes in gene expression as determined by Z-score, green is −2 and red is 2. (Middle) Representative top GO terms of upregulated and downregulated genes identified by differential expression analysis. (C) Venn diagrams showing number of unique or shared genes between ATAC- and RNA-seq datasets and charts for representative top gene ontology (GO) terms.

### Genome wide analysis for Dot1L effect on chromatin accessibility for thermogenic gene program

Thus far, we have shown that Dot1L-BKO mice have severely reduced thermogenic capacity due to significantly decreased BAT gene expression accompanied with reduced H3K79me2/3 at promoter regions of thermogenic genes. To study the underlying mechanism of how Dot1L ablation in BAT decreases H3K79me2/3 to decrease chromatin accessibility, thereby decreasing gene transcription, we used nuclei isolated from cold exposed Dot1L-BKO mice and their littermates to assess the chromatin landscape during thermogenesis by Assay for Transposase-Accessible Chromatin (ATAC-seq). Strikingly, Dot1L-BKO had significantly reduced peaks genome wide near the transcription start sites (TSS), while WT had concentrated open chromatin regions at TSS, as H3K79 methylation is reported as a gene activation marker. (Fig 5A left). With our primary interest in assessing Dot1L’s role in the regulation of thermogenesis, we compared relative ATAC-seq peaks for thermogenic genes, such as PRDM16, Zfp516, Cox8b and for mitochondria/oxidative phosphorylation, such as Cox7a1, Tfam, and Cyc1, in response to Dot1L ablation (Figure 5, middle). Also, Aifm2, Dot1L-BKO mice had much smaller peaks around promoters of UCP1, Cox8b and Elovl3 compared to WT mice (Figure 4, right), consistent with significantly decreased thermogenic gene expression in the Dot1L-BKO mice. Also, we found Dot1L-BKO had decreased expression of mitochondrial genes as well as Aifm2, a recently reported BAT-specific enzyme that converts NADH to NAD to sustain robust glycolysis to fuel thermogenesis (Nguyen et al., 2020). These data suggest that H3K79me2/3 by Dot1L is important for creating a chromatin landscape permissive to transcription in BAT upon cold stimulation.

Next, to examine gene expression changes upon Dot1L ablation in UCP1^+^ cells as a result of chromatin compaction, we performed RNA-seq using BAT from Dot1L-BKO mice and their littermates. Gene ontology of genes that were at least 2-fold downregulated in Dot1L-BKO mice revealed that mitochondrion organization, oxidative phosphorylation and cold-induced thermogenesis were some of the most affected processes (Figure 5B, middle). Notably, the downregulated thermogenic genes include UCP1, Cox8b, Elov3 and PGC1α as well as those related to mitochondrion organization and oxidative phosphorylation such as Nrf1, Cox20, Tomm13, and Immp21 (Figure 5B, left). With ATAC-seq results, these data suggest a mechanistic link between Dot1L-mediated chromatin remodeling and the global landscape of genome accessibility in brown fat for thermogenic gene expression. In addition, regulation of triglyceride and carbohydrate catabolism were compromised by Dot1L ablation. Conversely, chromosome organization, negative regulation of telomere maintenance, response to DNA damage were upregulated in the Dot1L knockout. In fact, we detected upregulated genes involved in chromatin compaction and gene repression, such as HDAC3, HDAC4 and DNMT1, further supporting that Dot1L increases chromatin accessibility (Figure 5B, right).

We then compared RNA-seq data to ATAC-seq obtained to identify common downregulated pathways found at lower levels using ATACs upon Dot1L ablation in UCP1+ cells, and we found about 40% of the total of downregulated genes were also detected from the ATAC-seq dataset (Figure 5C, left). Some of similar processes between the two datasets include regulation of cold-induced thermogenesis, transcription initiation/elongation factors, mitochondrial membrane organization as well as G-protein receptor signaling pathway (Figure 5C, right). Remarkably, the cluster of shared genes included thermogenic genes, such *as* UCP1, PGC1α, Cox8b, CideA, further underscoring the importance of Dot1L in brown adipose tissue. This suggest that Dot1L ablation results in reduced chromatin accessibility, leading to downregulated gene expression for those specific pathways. Altogether, we conclude that H3K79me2/3 by Dot1L plays a critical role in thermogenic gene activation by increasing chromatin accessibility.

## DISCUSSION

Numerous studies have supported the concept that a network of transcription factors and epigenetic coregulators must work together to fine-tune thermogenic program in response to environmental conditions (Sambeat et al., 2017). Here, we identify Dot1L, the only known H3K79 methyltransferase, as an interacting partner of Zc3h10, a transcriptional factor that activates UCP1, as well as Tfam and Nrf1, for thermogenesis. We show that Dot1L is recruited by Zc3h10 to the UCP1 and other thermogenic genes. We demonstrate that Dot1L directly interacts with Zc3h10 to co-occupy the same region of the UCP1 promoter. Zc3h10 binds to Dot1L via the 501-1000aa fragment, which is largely composed of coiled coil domains, evolutionarily conserved and widely-known for protein-protein interactions (Mier et al., 2017). This same coiled coil domain has been reported to interact with some of mixed lineage leukemia (MLL) oncogenic fusion proteins, such as AF10, to induce H3K79 methylation and constitutively activate a leukemic transcription program (Song et al., 2019). Here, we show that Dot1L is required for Zc3h10’s function for thermogenic gene program. In fact, Zc3h10 overexpression could not rescue the decreased thermogenic gene expression upon Dot1L ablation in mice, demonstrating that Dot1L is critical in Zc3h10 mediated activation of thermogenesis.

We establish that H3K79me2/3, catalyzed by Dot1L, as an activation mark for thermogenic gene program. In this regard, previous studies reported that H3K79 methylation is strongly associated with active transcription (Steger et al., 2008, Wood et al., 2018). Our Dot1L-BKO mice showed a markedly less open chromatin regions in promoter regions of thermogenic genes. The ATAC-seq data showed significantly reduced peaks genome-wide near the transcription start sites (TSS) in BAT of Dot1L-BKO mice, compared to control mice that had concentrated open chromatin regions at TSS upon cold exposure. In addition, studies reported that H3K79me2/3 may be active enhancer marks and may even be required for enhancer-promoter interactions to increase transcription (Markenscoff-Papadimitriou et al., 2014, Gilan et al., 2016, Godfrey et al., 2019). Godfrey et al. found H3K79me2/3 to be abundant at a subset of super-enhancers in MLL-related leukemia cells and that a loss of H3K79me2/3 by Dot1L inhibition led to significantly reduced enhancer-promoter interaction coupled with a decrease in transcription (Godfrey et al., 2019). We show that Dot1L is recruited by Zc3h10 to a UCP1 upstream region of -4.6 kb, a region suggested to be one of so-called super enhancers (SEs) (Whyte et al., 2013; Harms et al., 2015). In fact, Dot1L ablation in BAT of mice resulted in a significantly reduction of H3K79me2/3 at the -4.6 kb UCP1 region, correlating with decreased expression of UCP1. It is possible that H3K79me2/3 catalyzed by Dot1L may be important in maintaining enhancer/promoter association for transcriptional activation.

Due to limited accessibility of the H3K79 residue, located in the nucleosome core, how H3K79 methylation promotes gene activation is not clearly understood (van Leeuwen et al., 2002, Lu et al., 2008). Several studies support the concept that H3K79me2/3 and other well-profiled gene activation marks are interdependent (Steger et al., 2008). Steger et al. demonstrated that H3K79me2/3 at gene promoters highly correlates with H3K4me1/2/3 and H3K36me3, known gene activation marks in mammalian cells. Furthermore, Chen et al. reported that Dot1L may inhibit the localization of SIRT1 and SUV39H1, a H3K9ac demethylase and a H3K9me2 methyltransferase, respectively, at MLL fusion target genes correlating with elevated H3K9ac and low H3K9me2, to maintain an open chromatin state (Chen et al., 2015).

These authors proposed that H3K79me2/3 mark, in part, may function by inhibiting the histone deacetylase activity (Chen et al., 2015, Kang et al., 2018). In line with these findings, Dot1L probably works with other histone modifiers to maintain activation marks or to suppress repressive marks. Regardless, we establish that H3K79me2 and H3K79me3, in particular, function as activation marks for thermogenic gene program.

Overall, we demonstrate that Dot1L is critical for thermogenic program through H3K79 methylation and the requirement of Dot1L-Zc3h10 interaction for brown adipose thermogenesis in vitro and in vivo. With combined ATAC-seq and RNA-seq data, we provide molecular evidence that the decreased thermogenic gene expression is due to decreased H3K79me2/3 on promoter regions of thermogenic genes, as well as genes involved in chromatin remodeling.

Consequently, our mouse line of Dot1L ablated in UCP1^+^ cells show significantly decreased expression of thermogenic genes, including UCP1, resulting in cold intolerance with decreased oxygen consumption (Figure 4C-G), leading to increased adiposity as well as insulin insensitivity.

## MATERIALS AND METHODS

### Animals

Dot1L floxed mice (Dot1ltm1a(KOMP)Wtsi) were generated by the trans-NIH Knock-Out Mouse Project (KOMP) and obtained from the KOMP Repository (www.komp.org). These Dot1L floxed mice were first mated with FLPe mice from Jackson Lab. FLP mediated recombination was confirmed by PCR and resultant progeny were mated with UCP1-Cre mice (B6.FVB-Tg(Ucp1-cre)1Evdr/J) from Jackson Lab. Unless otherwise stated, male mice between 10 -14 weeks of age were used in experiments. Mice were fed a chow diet or a high fat diet (HFD) (45% fat derived calories-Dyets) ad libitum. EPZ-5676 was purchased from MedChemExpress (Cat. No.: HY-15593) and injected 20mg/kg/day for 8 days into BAT. All protocols for mice studies were approved from the University of California at Berkeley Animal Care and Use Committee.

### GST-pulldown

GST-Dot1L was purchased from epicypher. GST and GST-Dot1L plasmids were transformed into BL21(DE3) E. coli (NEB) and production was induced by 0.1M IPTG treatment. Resultant proteins were purified using Glutathione Sepharose 4B (GE) according to the manufacturer’s recommended protocol. [^35^S]-labeled Zc3h10 protein was produced by using TNT coupled transcription/translation kit (Promega). 20 ug of GST fusion proteins were incubated for 2 hrs at 4°C with in vitro translated Zc3h10 and glutathione sepharose beads. The beads were washed 3 times with binding buffer, and bound proteins were eluted by boiling in Laemmli sample buffer, separated by SDS-PAGE and analyzed by autoradiography.

### Tandem affinity purification (TAP) and mass spectrometry Analysis

10ug of Zc3h10-CTAP or CTAP vector plasmid were transfected in 293FT cells using lipofectamine 2000 (Invitrogen). Cells were lysed and immunoprecipated using buffers from the InterPlay Mammalian TAP system manufacturer recommended with the noted exception: after binding of the cell lysates to the Streptavidin resin, the resin was re-incubated with 200 ug of BAT nuclear extracts diluted to 1 mL in SBB overnight. TAP eluates were boiled in SDS, ran on an SDS-PAGE gel, and stained with Coomassie Brilliant Blue. Bands that were identified to be specific to the Zc3h10-TAP lane were excised from both the Zc3h10-TAP and TAP vector lanes and proteins were identified by mass spectrometry performed by the Vincent J. Coates Proteomics/ Mass Spectrometry Laboratory (P/MSL) at UC Berkeley.

### Cold-induced Thermogenesis

Core body temperature was determined using a Physitemp BAT-12 probe at 4°C.

### Indirect Calorimetry

Oxygen Consumption was measured using the Comprehensive Laboratory Animal Monitoring System (CLAMS). Data were normalized to lean body mass determined by EchoMRI. Mice were individually caged and maintained under a 12 hr light/12 hr dark cycle. Food consumption and locomotor activity were tracked.

### GTT and ITT

For GTTs, mice were fasted overnight, and glucose (2 mg/g) was administered intraperitoneally. For ITTs, mice were fasted 4 hr, and insulin (0.75 U/kg) was administered.

### Cell culture

HEK293FT and 3T3-L1 cells were obtained from UCB Cell Culture Facility supported by The University of California Berkeley. The immortalized BAT cell line was from Dr. Shingo Kajimura (Harvard). Cells were grown in standard condition with 5% CO_2_, at 37°C. BAT cells, 3T3-L1 cells and 293FT cells were maintained in DMEM containing 10% FBS and 1% pen/strep prior to differentiation/ transfection. Brown adipocyte differentiation was performed as described in (Yi et al., 2019). Ad-m-Dot1L-shRNA, Ad-Zc3h10-flag adenovirus and Ad-GFP-m-DOT1L were purchased from Vector Biolabs. Knockdown of Dot1L accomplished using adenoviral transduction of MOI of 250 at day 4 of brown adipocyte differentiation. Differentiation of 3T3-L1 cells was induced by treating confluent cells with DMEM containing 10% FBS, 850 nM insulin, 0.5mM isobutyl-methylxanthine, 1 µM dexamethasone, 1 nM T3, 125 nM indomethacin. After 48 hrs of induction, cells were switched to a maintenance medium containing 10% FBS, 850 nM insulin and 1 nM T3. 3T3-L1 cells were infected on day 4 of differentiation using either GFP, Dot1L or Zc3h10 adenovirus. Viral medium was replaced by maintenance medium the following day. To stimulate thermogenesis, differentiated 3T3-L1 cells were treated 6 hrs with 10 µM forskolin on day 6. Inhibition of Dot1L was accomplished by addition of either 5 nM EPZ5676 (MedChemExpress, Cat. No.: HY-15593) or DMSO.

Inducible Lentiviral Nuclease hEF1α-Blast-Cas9 (GE Healthcare) was packaged into lentivirus by using MISSION Lentiviral Packaging Mix (Sigma). BAT inducible Cas9 cell line was then generated by transduced BAT cells with inducible Cas9 lentivirus and then selected with blasticidin (10μg/ml). Two stable Dot1L sgRNA expressing cell lines were generated by transducing in inducible Cas9 cells with lentivirus containing two sgRNA with target sequences of GTCTCGTGCAGCATAACCAG, or ACGCCGTGTTGTATGCATCT (Abmgood) and were then selected by neomycin (400μg/ml). These Dot1L KO pools were subjected to brown adipocyte differentiation. After 48 hr of induction, cells were treated with doxycycline (1ug/ml).

### Oxygen consumption measurement

BAT cells were differentiated in 12-well plates, trysinized, and reseeded in XF24 plates at 50 K cells per well at day 4 of differentiation and assayed on day 5 of differentiation. On the day of experiments, the cells were washed 3 times and maintained in XF-DMEM (Sigma-Aldrich) supplemented with 1 mM sodium pyruvate and 17.5 mM glucose. Oxygen consumption was blocked by 1 μM oligomycin. Maximal respiratory capacity was assayed by the addition of 1 μM FCCP. Tissues were incubated for 1 hr at 37°C without CO_2_ prior to analysis on the Seahorse XF24 Analyzer. Uncoupled respiration was calculated as OCR under oligomycin treatment minus OCR under antimycin A/rotenone treatment.

### RT-qPCR Analysis and Western blotting

Reverse transcription was performed with 500 ng of total RNA using SuperScript III (Invitrogen). RT-qPCR was performed in triplicate with an ABI PRISM 7500 sequence detection system (Applied Biosystems) to quantify the relative mRNA levels for various genes. Statistical analysis of the qPCR was obtained using the ΔΔCt (2^-ddCT) method with Eef1a1 or 18s as the control. RT-qPCR primer sets are listed in Table 1. Library generation for RNA sequencing was carried out at the Functional Genomics Laboratory at UC Berkeley.

**Table 1:**
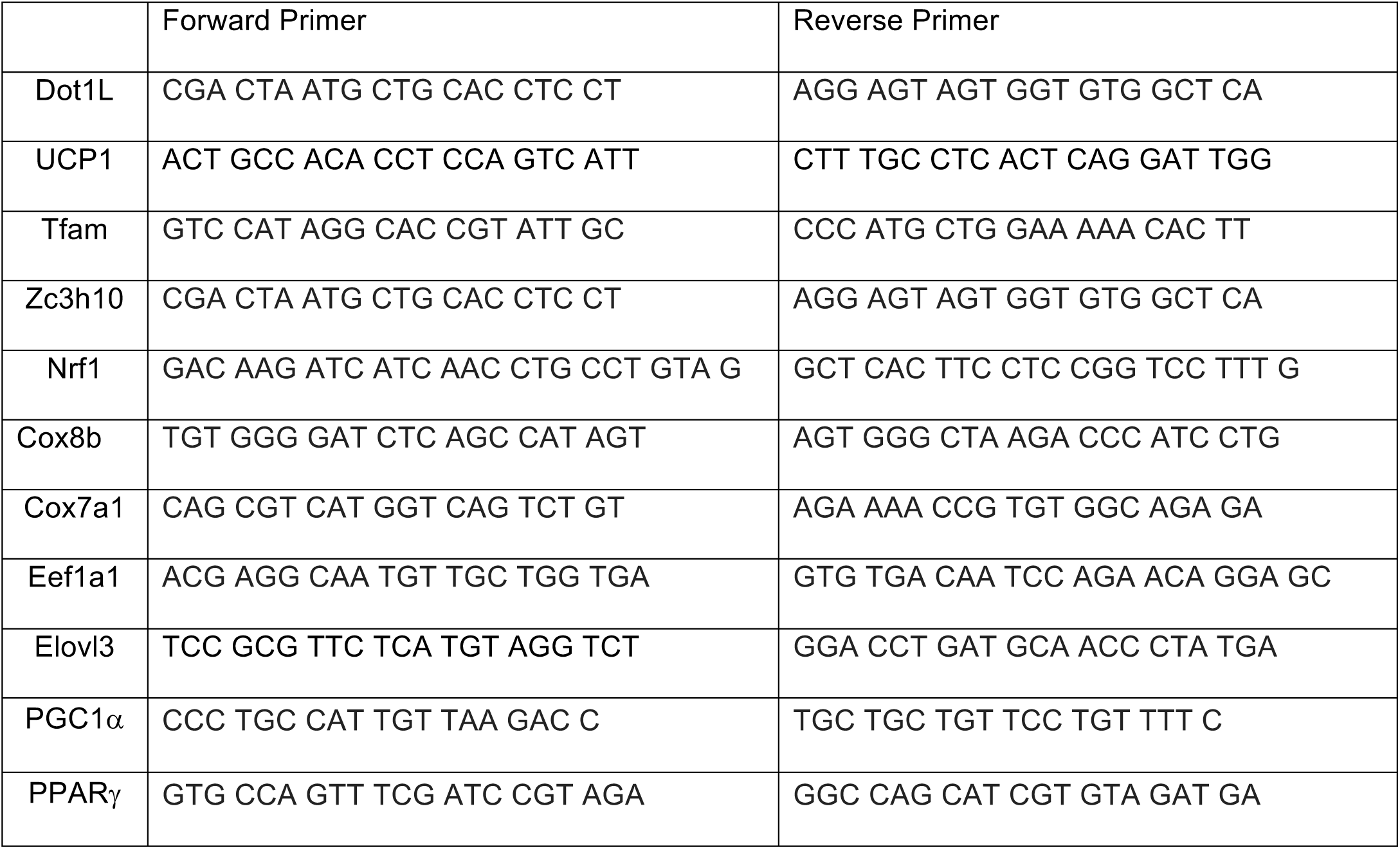
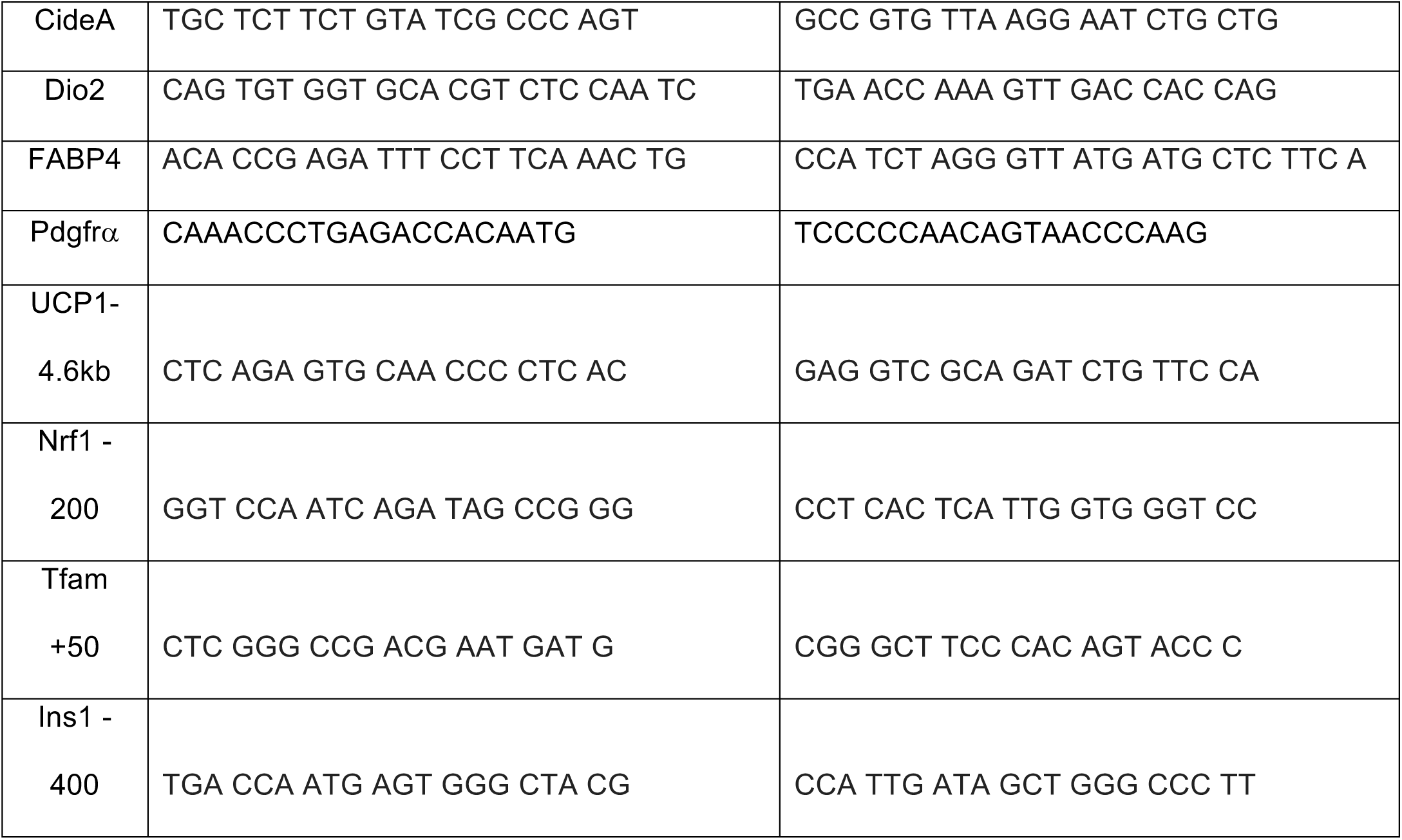
Primer Sets used for RT-qPCR.

For western blot analysis, total cell lysates were prepared using RIPA buffer and nuclear extracts were prepared using the NE-PER Nuclear and Cytoplasmic Extraction kit (Thermo). Proteins were separated by SDS-PAGE, transferred to nitrocellulose membrane and probed with the indicated antibodies.

### Antibodies

The following antibodies were used where indicated: UCP1 (Abcam, 10983), Zc3h10 (Thermo Fisher, PA5-31814; for immunoblotting), Zc3h10 (Abnova, H00084872-B01P; for ChIP), GAPDH (Cell signaling, 5174S), Dot1L (Bethyl, A300-953A), FLAG (Cell signaling, 14793S), HA (Cell signaling, 3724S), Tubulin (Abcam, Ab52866), Histone H3 (Cell signaling, 4499S), H3K79me3 (Abcam, ab2621), H3K79me2 (Abcam, ab3594)

### Luciferase-reporter Assay

293FT cells were transfected with 300 ng Zc3h10 and/or Dot1L expression plasmid, together with 100 ng of indicated luciferase reporter construct and 0.5 ng pRL-CMV in 24-well plates. Cells were lysed 48 hr post-transfection and assayed for luciferase activity using the Dual-Luciferase Kit (Promega) according to the manufacturer’s recommended protocol.

### Plasmid Constructs

The HA-Dot1L expression vector was purchased from Genecopoeia. The Dot1L sequence was subcloned by PCR amplifying and inserting into CTAP vector from Agilent, as well as pGEX-4T-3 from GE.

### Coimmunoprecipitation

For Co-IP experiments using tagged constructs, 293FT cells were transfected using Lipofectamine 2000 to express FLAG-tagged Zc3h10 and HA-tagged Dot1L. Cells were lysed in IP buffer containing 20 mM Tris, pH 7.4, 150 mM NaCl, 1 mM EDTA, 10% glycerol, 1% NP-40 supplemented with protease inhibitors. Total cell lysates were incubated 2 hrs at 4°C with either anti-FLAG M2 for Zc3h10 or anti-HA for Dot1L. For Co-IP in brown adipose tissue of wild type mice, nuclear extraction was carried out using the NE-PER Nuclear and Cytoplasmic Extraction kit (Thermo). Equal amounts of nuclear extracts were incubated with specific antibodies and protein A/G agarose beads overnight at 4°C. Agarose beads were washed 3 times and bound proteins were eluted by boiling in Laemmli sample buffer and analysed by immunoblotting using the indicated antibodies.

### ChIP-qPCR

ChIP was performed using ChIP kit (Cell signaling, 57976s). Briefly, BAT cells overexpressing Dot1L or Zc3h10 were fixed with disuccinimidyl glutarate (DSG) at 2 mM concentration in PBS for 45 min at room temperature before cross-linking with 1% methanol-free formaldehyde in PBS for 10 min. For ChIP experiments using BAT, tissues from 3 BATs were combined and minced on ice. The reaction was stopped by incubating with 125 mM glycine for 10 min. Cells or tissues were rinsed with ice-cold phosphate-buffered saline (PBS) for 3 times, and lysed in IP lysis buffer containing 500 mM HEPES-KOH, pH 8, 1 mM EDTA, 0.5 mM EGTA, 140 mM NaCl, 0.5% NP-40, 0.25% Triton X-100, 10% glycerol, and protease inhibitors for 10 min at 4 °C. Nuclei were collected by centrifugation at 600 × g for 5 min at 4 °C. Nuclei were released by douncing on ice and collected by centrifugation. Nuclei were then lysed in nuclei lysis buffer containing 50 mM Tris, pH 8.0, 1% SDS 10 mM EDTA supplemented with protease inhibitors, and sonicated 3 times by 20 s burst, each followed by 1 min cooling on ice. Chromatin samples were diluted 1:10 with the dilution buffer containing 16.7 mM Tris pH 8.1, 0.01% SDS 1.1 % Triton X-100 1.2 mM EDTA, 1.67 mM NaCl, and proteinase inhibitor cocktail. Soluble chromatin was quantified by absorbance at 260 nm, and equivalent amounts of input DNA were immunoprecipitated using 10 μg of indicated antibodies or normal mouse IgG (Santa Cruz) and protein A/G magnetic beads (Thermo-Fisher). After the beads were washed and cross-linking reversed, DNA fragments were purified using the Simple ChIP kit (cell signaling). Samples were analyzed by qPCR using the primer sets in Table 1 for enrichment in target areas. The promoter occupancy for UCP1 and other indicated genes was confirmed by RT-qPCR. The fold enrichment values were normalized to input.

### RNA-seq

WT and Dot1L-BKO mice were cold exposed for 2 hrs. Total RNA from BAT was prepared using RNEasy kit (Qiagen). Strand-specific libraries were generated from 500 ng total RNA using the TruSeq Stranded Total RNA Library Prep Kit (Illumina). cDNA libraries were single-end sequenced (50 bp) on an Illumina HiSeq 2000 or 4000. Using Partek software, reads were aligned to the mouse genome (NCBI38/mm10). A gene was included in the analysis if it met all the following criteria: the maximum RPKM reached 4 unit at any time point, the gene length was >100 bp. Selected genes were induced at least +/-1.8-fold, and the expression was significantly different from the basal (p<0.05).

### ATAC-seq

For ATAC-seq, we used two mice per condition and processed nuclei extracted from BAT separately, thus each sample analyzed was a separate biological replicate. ATAC-seq was performed using Nextera DNA library Preparation kit (Illumina, 15028212) according to (Chen et al., 2018). In all, 5 × 10^4^ nuclei from tissues were collected in lysis buffer containing 10 mM Tris pH 7.4, 10 mM NaCl, 3 mM MgCl2, and 1% NP-40, and spun at 500 × g at 4 °C for 10 min. The pellets were resuspended in the transposase reaction mixture containing 25 μL 2 × Tagmentation buffer, 2.5 μL transposase, and 22.5 μL nuclease-free water, and incubated at 37 °C for 30 min. The samples were purified using MinElute PCR Purification kit (Qiagen, 28006) and amplification was performed in 1 × next PCR master mix (NEB, M0541S) and 1.25 μM of custom Nextera PCR primers 1 and 2 with the following PCR conditions: 72 °C for 5 min; 98 °C for 30 s; and thermocycling at 98 °C for 10 s, 63 °C for 30 s, and 72 °C for 1 min.

Samples were amplified for five cycles and 5 μL of the PCR reaction was used to determine the required cycles of amplification by qPCR. The remaining 45 μL reaction was amplified with the determined cycles and purified with MinElute PCR Purification kit (Qiagen, 28006) yielding a final library concentration of ∼30 nM in 20 μL. Libraries were subjected to pair-end 50 bp sequencing on HiSeq4000 with 4–6 indexed libraries per lane. Partek Flow Genomics Suite was used to analyze sequencing data. Reads were aligned to mm10 genome assembly using BWA-backtrack. MACS2 in ATAC mode was then used to identify peaks from the aligned reads with a q-value cutoff of 0.05 and fold enrichment cutoff of 2.0. Quantify regions tool was used to quantify the peaks identified by MACS2 from each sample to generate a union set of regions.

Regions were then annotated to RefSeq mRNA database and analysis of variance (ANOVA) was performed to determine significant peak differences between groups. UCSC genome browser was used to visualize peaks. Gene set enrichment and pathway analysis was used to highlight biological processes with the highest q-values among those identified.

### Statistical analysis

Statistical comparisons were made using a two-tailed unpaired t-test using GraphPad Prism 8 software (GraphPad Software Inc., La Jolla, CA, USA). For genome-wide analyses, we employed Partek Genomics Suite (Partek Inc., St. Louis, Missouri, USA) using ANOVA for ATAC-seq comparisons, and DESeq2 for RNA-seq differential expression comparisons, and subsequently used Gene set enrichment tool for gene ontology to identify the significantly affected pathways. Data are expressed as means ± standard errors of the means (SEM). The statistical differences in mean values were assessed by Student’s *t* test. All experiments were performed at least twice and representative data are shown.

## AUTHOR CONTRIBUTIONS

D.Y. performed loss-of-function/gain-of-function studies in vitro/in vivo experiments. H.P.N. performed the interaction studies, assisted in animal studies and edited the manuscript. J.D. assisted in animal studies. J.V. made plasmid constructs and performed bioinformatics analysis and edited the manuscript. Y.X. assisted in RT-qPCR/immunoblotting. J.M.D. found and characterized Dot1L. Y.W. performed ChIP and ATAC-seq. H.S.S. designed the project and guided experiments. D.Y. and H.S.S wrote the manuscript.

## ACKLOWLDGEMENTS

We thank M. Zhu for technical assistance. The work was supported by NIH grant DK120075 to H.S.S.

## SUPPLEMENTARY MATERIALS

**S1.**
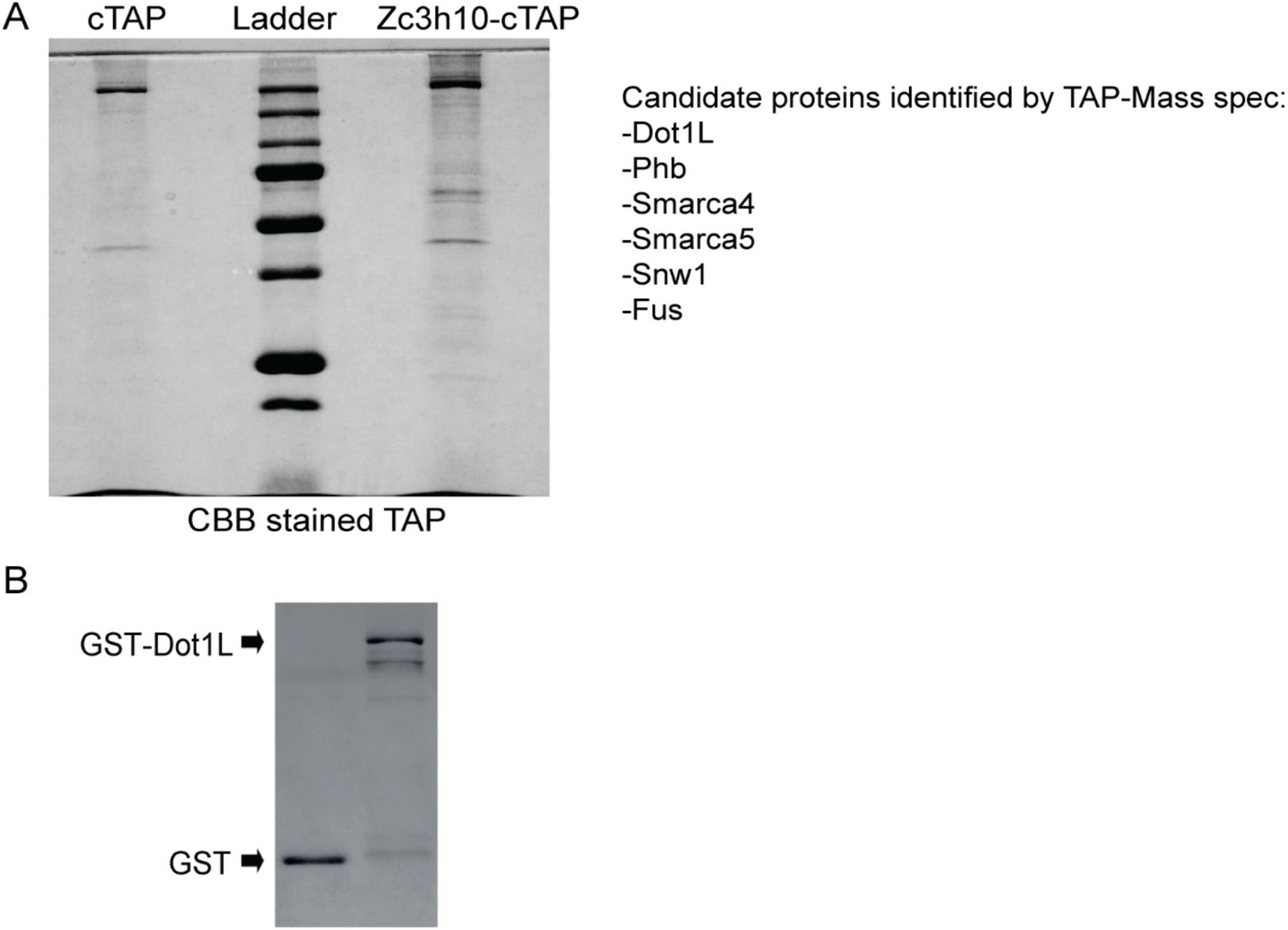
DotIL directly interacts with Zc3h10 for UCP1 promoter activation. (A) Gel stained with Coomassie Blue after cTAP purification assay and a list of potential Zc3h10 interacting proteins identified by Mass spec. (B) Coomassie staining for GST alone and GST-Dot1 L fusion protein.

**S2.**
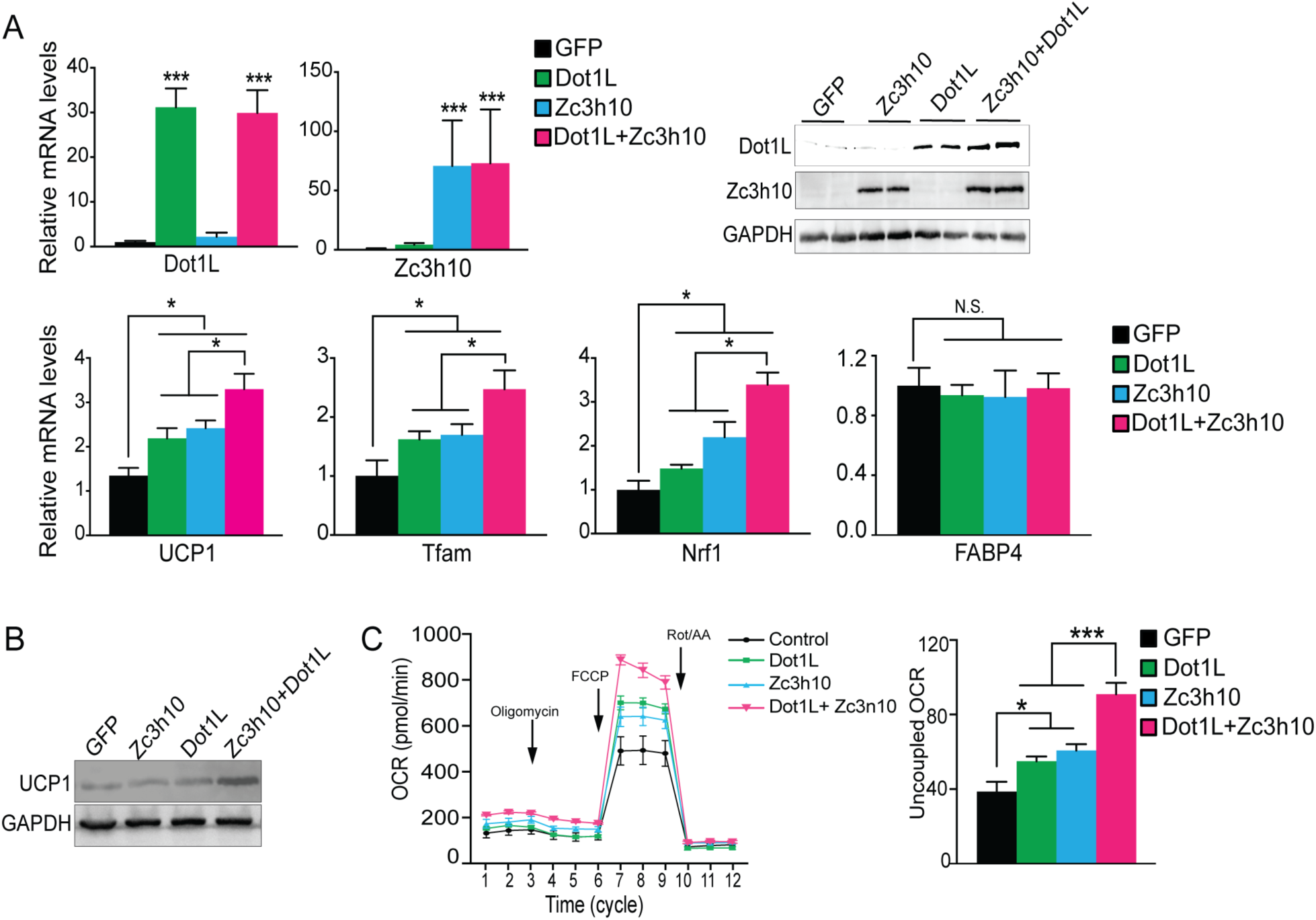
Dot1L promotes the thermogenic gene program in vitro. (A) RT-qPCR for indicated genes and immunoblotting for indicated proteins in 3T3-L1 cells that were transduced with either AdGFP or AdZc3h10 or AdDotIL individually or in combination for overexpression (OE) of Zc3h10 and DotIL (n=6). The differentiated cells were treated with forskolin (10uM) for 6 hr to induce beiging. (B) Immunoblotting for UCP1. (C) (Left) OCR measured in Zc3h10 OE and DotlLOE, individually and in combination in differentiated 3T3-L1 adipocytes that were induced to beiging (n=6). (Right) Uncoupled OCR under oligomycin (0.5 uM).

**S3.**
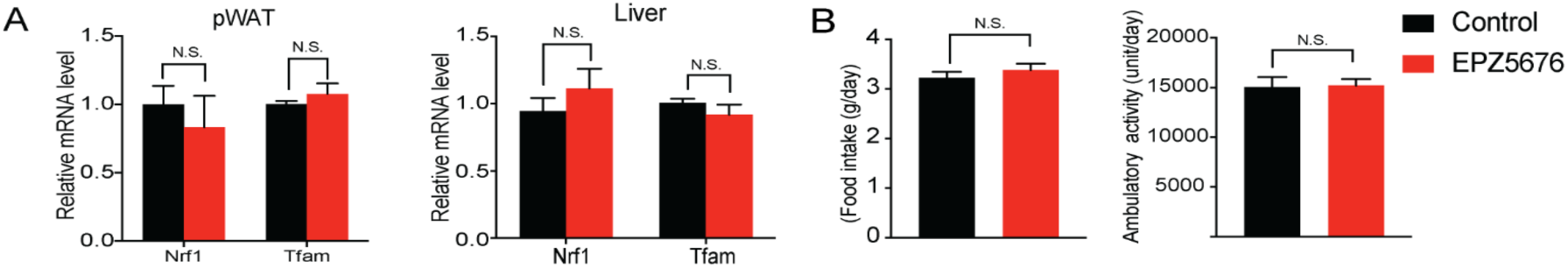
Inhibition of DotIL activity impairs BAT gene program in vivo. (A) RT-qPCR for indicated genes in pWAT and liver of control and EPZ5676 injected mice. (B) Food intake and locomotor activity of control and EPZ5676 injected mice.

**S4.**
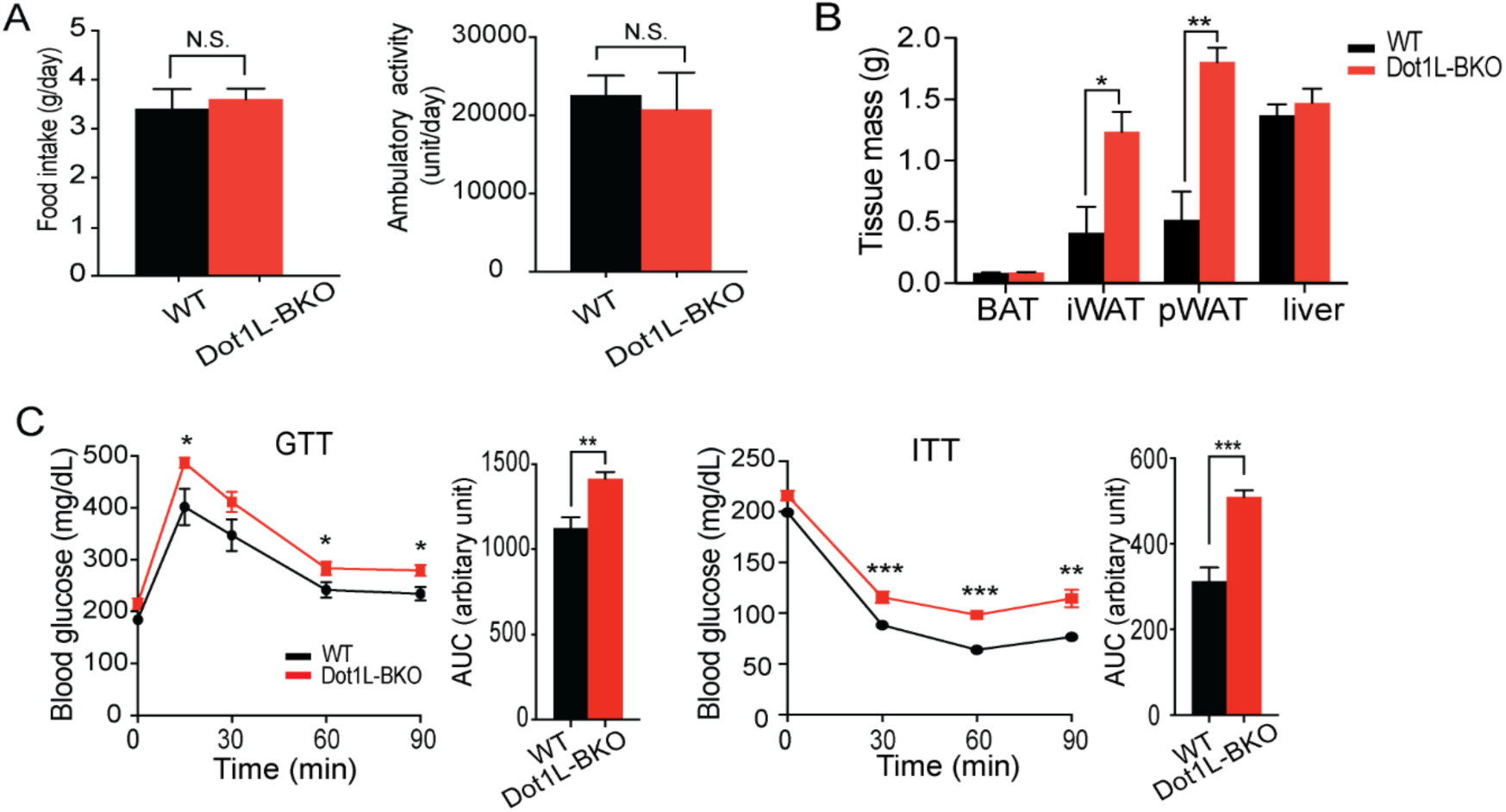
DotIL is required for cold-induced thermogenesis in mice. (A) Food intake and locomotor activity of WT and Dot1L-BKO mice. (B) Body weight and mass of various tissues taken from WT and Dot1L-BKO mice at 14-week of age. (C) GTT and ITT of Dot1 L-BKO mice at 14 wk of age that were on HFD for 3 weeks (n=5).

## REFERENCES

Cannon, B. & Nedergaard, J. 2004. Brown adipose tissue: function and physiological significance. Physiol Rev, 84, 277–359.

Chen, C. W., Koche, R. P., Sinha, A. U., Deshpande, A. J., Zhu, N., Eng, R., Doench, J. G., Xu, H., Chu, S. H., Qi, J., Wang, X., Delaney, C., Bernt, K. M., Root, D. E., Hahn, W. C., Bradner, J. E. & Armstrong, S. A. 2015. DOT1L inhibits SIRT1-mediated epigenetic silencing to maintain leukemic gene expression in MLL-rearranged leukemia. Nat Med, 21, 335–43.

Chen, D., Liu, W., Zimmerman, J., Pastor, W. A., Kim, R., Hosohama, L., Ho, J., Aslanyan, M., Gell, J. J., Jacobsen, S. E. & Clark, A. T. 2018. The TFAP2C-Regulated OCT4 Naive Enhancer Is Involved in Human Germline Formation. Cell Rep, 25, 3591–3602 e5.

Chondronikola, M., Volpi, E., Borsheim, E., Porter, C., Annamalai, P., Enerback, S., Lidell, M. E., Saraf, M. K., Labbe, S. M., Hurren, N. M., Yfanti, C., Chao, T., Andersen, C. R., Cesani, F., Hawkins, H. & Sidossis, L. S. 2014. Brown adipose tissue improves whole-body glucose homeostasis and insulin sensitivity in humans. Diabetes, 63, 4089–99.

Cypess, A. M., Lehman, S., Williams, G., Tal, I., Rodman, D., Goldfine, A. B., Kuo, F. C., Palmer, E. L., Tseng, Y., Doria, A., Kolodny, G. M. & Kahn, C. R. 2009. Identification and Importance of Brown Adipose Tissue in Adult Humans. New England Journal of Medicine, 360, 1509–1517.

Dempersmier, J., Sambeat, A., Gulyaeva, O., Paul, S. M., Hudak, C. S., Raposo, H. F., Kwan, H. Y., Kang, C., Wong, R. H. & Sul, H. S. 2015. Cold-inducible Zfp516 activates UCP1 transcription to promote browning of white fat and development of brown fat. Mol Cell, 57, 235–46.

Farmer, S. R. 2008. Molecular determinants of brown adipocyte formation and function. Genes Dev, 22, 1269–75.

Feng, Q., Wang, H., Ng, H. H., Erdjument-Bromage, H., Tempst, P., Struhl, K. & Zhang, Y. 2002. Methylation of H3-lysine 79 is mediated by a new family of HMTases without a SET domain. Curr Biol, 12, 1052–8.

Frederiks, F., Tzouros, M., Oudgenoeg, G., Van Welsem, T., Fornerod, M., Krijgsveld, J. & Van Leeuwen, F. 2008. Nonprocessive methylation by Dot1 leads to functional redundancy of histone H3K79 methylation states. Nat Struct Mol Biol, 15, 550–7.

Gilan, O., Lam, E. Y., Becher, I., Lugo, D., Cannizzaro, E., Joberty, G., Ward, A., Wiese, M., Fong, C. Y., Ftouni, S., Tyler, D., Stanley, K., Macpherson, L., Weng, C. F., Chan, Y. C., Ghisi, M., Smil, D., Carpenter, C., Brown, P., Garton, N., Blewitt, M. E., Bannister, A. J., Kouzarides, T., Huntly, B. J., Johnstone, R. W., Drewes, G., Dawson, S. J., Arrowsmith, C. H., Grandi, P., Prinjha, R. K. & Dawson, M. A. 2016. Functional interdependence of BRD4 and DOT1L in MLL leukemia. Nat Struct Mol Biol, 23, 673–81.

Godfrey, L., Crump, N. T., Thorne, R., Lau, I. J., Repapi, E., Dimou, D., Smith, A. L., Harman, J. R., Telenius, J. M., Oudelaar, A. M., Downes, D. J., Vyas, P., Hughes, J. R. & Milne, T. A. 2019. DOT1L inhibition reveals a distinct subset of enhancers dependent on H3K79 methylation. Nat Commun, 10, 2803.

Hibi, M., Oishi, S., Matsushita, M., Yoneshiro, T., Yamaguchi, T., Usui, C., Yasunaga, K., Katsuragi, Y., Kubota, K., Tanaka, S. & Saito, M. 2016. Brown adipose tissue is involved in diet-induced thermogenesis and whole-body fat utilization in healthy humans. International Journal of Obesity, 40, 1655–1661.

Jimenez, M. A., Akerblad, P., Sigvardsson, M. & Rosen, E. D. 2007. Critical role for Ebf1 and Ebf2 in the adipogenic transcriptional cascade. Mol Cell Biol, 27, 743–57.

Kang, J. Y., Kim, J. Y., Kim, K. B., Park, J. W., Cho, H., Hahm, J. Y., Chae, Y. C., Kim, D., Kook, H., Rhee, S., Ha, N. C. & Seo, S. B. 2018. KDM2B is a histone H3K79 demethylase and induces transcriptional repression via sirtuin-1-mediated chromatin silencing. FASEB J, 32, 5737–5750.

Kong, X., Banks, A., Liu, T., Kazak, L., Rao, R. R., Cohen, P., Wang, X., Yu, S., Lo, J. C., Tseng, Y. H., Cypess, A. M., Xue, R., Kleiner, S., Kang, S., Spiegelman, B. M. & Rosen, E. D. 2014. IRF4 is a key thermogenic transcriptional partner of PGC-1alpha. Cell, 158, 69–83.

Kriszt, R., Arai, S., Itoh, H., Lee, M. H., Goralczyk, A. G., Ang, X. M., Cypess, A. M., White, A. P., Shamsi, F., Xue, R., Lee, J. Y., Lee, S. C., Hou, Y., Kitaguchi, T., Sudhaharan, T., Ishiwata, S., Lane, E. B., Chang, Y. T., Tseng, Y. H., Suzuki, M. & Raghunath, M. 2017. Optical visualisation of thermogenesis in stimulated single-cell brown adipocytes. Sci Rep, 7, 1383.

Lichtenbelt, W. D. V., Vanhommerig, J. W., Smulders, N. M., Drossaerts, J. M. A. F. L., Kemerink, G. J., Bouvy, N. D., Schrauwen, P. & Teule, G. J. J. 2009. Cold-Activated Brown Adipose Tissue in Healthy Men (vol 360, pg 1500, 2009). New England Journal of Medicine, 360, 1917–1917.

Lu, X., Simon, M. D., Chodaparambil, J. V., Hansen, J. C., Shokat, K. M. & Luger, K. 2008. The effect of H3K79 dimethylation and H4K20 trimethylation on nucleosome and chromatin structure. Nat Struct Mol Biol, 15, 1122–4.

Markenscoff-Papadimitriou, E., Allen, W. E., Colquitt, B. M., Goh, T., Murphy, K. K., Monahan, K., Mosley, C. P., Ahituv, N. & Lomvardas, S. 2014. Enhancer interaction networks as a means for singular olfactory receptor expression. Cell, 159, 543–57.

Mier, P., Alanis-Lobato, G. & Andrade-Navarro, M. A. 2017. Protein-protein interactions can be predicted using coiled coil co-evolution patterns. J Theor Biol, 412, 198–203.

Min, J., Feng, Q., Li, Z., Zhang, Y. & Xu, R. M. 2003. Structure of the catalytic domain of human DOT1L, a non-SET domain nucleosomal histone methyltransferase. Cell, 112, 711–23.

Ng, H. H., Feng, Q., Wang, H., Erdjument-Bromage, H., Tempst, P., Zhang, Y. & Struhl, K. 2002. Lysine methylation within the globular domain of histone H3 by Dot1 is important for telomeric silencing and Sir protein association. Genes Dev, 16, 1518–27.

Nguyen, A. T. & Zhang, Y. 2011. The diverse functions of Dot1 and H3K79 methylation. Genes Dev, 25, 1345–58.

Nguyen, H. P., Yi, D., Lin, F., Viscarra, J. A., Tabuchi, C., Ngo, K., Shin, G. W., Lee, A. Y. F., Wang, Y. H. & Sul, H. S. 2020. Aifm2, a NADH Oxidase, Supports Robust Glycolysis and Is Required for Cold- and Diet-Induced Thermogenesis. Molecular Cell, 77, 600-+.

Okada, Y., Feng, Q., Lin, Y., Jiang, Q., Li, Y., Coffield, V. M., Su, L., Xu, G. & Zhang, Y. 2005. hDOT1L links histone methylation to leukemogenesis. Cell, 121, 167–78.

Puigserver, P., Wu, Z., Park, C. W., Graves, R., Wright, M. & Spiegelman, B. M. 1998. A cold-inducible coactivator of nuclear receptors linked to adaptive thermogenesis. Cell, 92, 829–39.

Rajakumari, S., Wu, J., Ishibashi, J., Lim, H. W., Giang, A. H., Won, K. J., Reed, R. R. & Seale, P. 2013. EBF2 determines and maintains brown adipocyte identity. Cell Metab, 17, 562–74.

Sambeat, A., Gulyaeva, O., Dempersmier, J. & Sul, H. S. 2017. Epigenetic Regulation of the Thermogenic Adipose Program. Trends Endocrinol Metab, 28, 19–31.

Sambeat, A., Gulyaeva, O., Dempersmier, J., Tharp, K. M., Stahl, A., Paul, S. M. & Sul, H. S. 2016. LSD1 Interacts with Zfp516 to Promote UCP1 Transcription and Brown Fat Program. Cell Rep, 15, 2536–49.

Sanchez-Gurmaches, J., Hung, C. M. & Guertin, D. A. 2016. Emerging Complexities in Adipocyte Origins and Identity. Trends in Cell Biology, 26, 313–326.

Schulze, J. M., Jackson, J., Nakanishi, S., Gardner, J. M., Hentrich, T., Haug, J., Johnston, M., Jaspersen, S. L., Kobor, M. S. & Shilatifard, A. 2009. Linking cell cycle to histone modifications: SBF and H2B monoubiquitination machinery and cell-cycle regulation of H3K79 dimethylation. Mol Cell, 35, 626–41.

Seale, P., Bjork, B., Yang, W., Kajimura, S., Chin, S., Kuang, S., Scime, A., Devarakonda, S., Conroe, H. M., Erdjument-Bromage, H., Tempst, P., Rudnicki, M. A., Beier, D. R. & Spiegelman, B. M. 2008. PRDM16 controls a brown fat/skeletal muscle switch. Nature, 454, 961–7.

Song, X., Yang, L., Wang, M., Gu, Y., Ye, B., Fan, Z., Xu, R. M. & Yang, N. 2019. A higher-order configuration of the heterodimeric DOT1L-AF10 coiled-coil domains potentiates their leukemogenenic activity. Proc Natl Acad Sci U S A, 116, 19917–19923.

Steger, D. J., Lefterova, M. I., Ying, L., Stonestrom, A. J., Schupp, M., Zhuo, D., Vakoc, A. L., Kim, J. E., Chen, J., Lazar, M. A., Blobel, G. A. & Vakoc, C. R. 2008. DOT1L/KMT4 recruitment and H3K79 methylation are ubiquitously coupled with gene transcription in mammalian cells. Mol Cell Biol, 28, 2825–39.

Van Leeuwen, F., Gafken, P. R. & Gottschling, D. E. 2002. Dot1p modulates silencing in yeast by methylation of the nucleosome core. Cell, 109, 745–756.

Virtanen, K. A., Lidell, M. E., Orava, J., Heglind, M., Westergren, R., Niemi, T., Taittonen, M., Laine, J., Savisto, N. J., Enerback, S. & Nuutila, P. 2009. Functional brown adipose tissue in healthy adults. N Engl J Med, 360, 1518–25.

Wang, W. S. & Seale, P. 2016. Control of brown and beige fat development. Nature Reviews Molecular Cell Biology, 17, 691–702.

Wood, K., Tellier, M. & Murphy, S. 2018. DOT1L and H3K79 Methylation in Transcription and Genomic Stability. Biomolecules, 8.

Yi, D., Dempersmier, J. M., Nguyen, H. P., Viscarra, J. A., Dinh, J., Tabuchi, C., Wang, Y. & Sul, H. S. 2019. Zc3h10 Acts as a Transcription Factor and Is Phosphorylated to Activate the Thermogenic Program. Cell Rep, 29, 2621–2633 e4.

Yi, D., Nguyen, H. P. & Sul, H. S. 2020. Epigenetic dynamics of the thermogenic gene program of adipocytes. Biochem J, 477, 1137–1148.

